# Deep Phosphoproteomic Elucidation of Metformin-Signaling in Heterogenous Colorectal Cancer Cells

**DOI:** 10.1101/2022.07.07.499038

**Authors:** Barbora Salovska, Erli Gao, Sophia Müller-Dott, Wenxue Li, Aurelien Dugourd, George Rosenberger, Julio Saez-Rodriguez, Yansheng Liu

## Abstract

The biguanide drug metformin is a safe and widely prescribed drug for type 2 diabetes. Interestingly, hundreds of clinical trials were set to evaluate the potential role of metformin in the prevention and treatment of cancer including colorectal cancer (CRC). To interrogate cell signaling events and networks in CRC and explore the druggability of the metformin-rewired phosphorylation network, we performed a proteomic and phosphoproteomic analysis on a panel of 12 molecularly heterogeneous CRC cell lines. Using in-depth data-independent analysis mass spectrometry (DIA-MS), we profiled a total of 10,142 proteins and 56,080 phosphosites (P-sites) in CRC cells treated with metformin for 30 minutes and 24 hours. Our results indicate that metformin does not directly trigger or inhibit any immediate phosphorylation events. Instead, it primarily remodels cell signaling in the long-term. Strikingly, the phosphorylation response to metformin was highly heterogeneous in the CRC panel, uncovering four groups of metformin responsivity. We further performed a network analysis to systematically estimate kinase/phosphatase activities and reconstruct signaling cascades in each cell line. We created a “MetScore” which catalogs the most consistently perturbed P-sites among CRC cells for future studies. Finally, we leveraged the metformin P-site signature to identify pharmacodynamic interactions revealing a number of candidate metformin-interacting drugs. Together, we provide a data resource using state-of-the-art phosphoproteomics to understand the metformin-induced cell signaling for potential cancer therapeutics.

## Introduction

The biguanide drug metformin is the first-line treatment for type 2 diabetes (T2D) that has been used in Europe since 1960s and was approved by the Food and Drug Administration (FDA) in the 1990s (Bailey and Turner, 1996). The drug has a superior safety profile and is highly tolerated with a minimum of serious side-effects compared to other biguanides (Lalau and Race, 2001) and has been used by more than 150 millions of people worldwide (He and Wondisford, 2015). In addition to the antihyperglycemic effect, accumulating evidence has suggested multiple beneficial effects of metformin therapy including weight loss in obese patients, cardiorenal protection, neuroprotection, and overall health improvement (Kulkarni et al., 2020). Consequently, metformin has been used off-label to support the treatment of other conditions such as prediabetes, type 1 diabetes, gestational diabetes, polycystic ovary syndrome, diabetic nephropathy, or non-alcoholic fatty liver disease (Drzewoski and Hanefeld, 2021). Moreover, a retrospective study published in 2005 reported for the first time that metformin reduced cancer risk in diabetic patients (Evans et al., 2005). Since then, metformin has been investigated in 400 clinical trials (June 2022, ClinicalTrials.gov) for its potential in preventing or treatment of various types of cancer.

Colorectal cancer (CRC) has been one of the most prevalent cancer types and one of the leading causes of cancer death globally (Sung et al., 2021). Moreover, a meta-analysis systematically investigating the association between CRC and T2D revealed a 30% increased risk of developing CRC in diabetic patients (Godsland, 2009; Larsson et al., 2005). Mounting evidence indicates a potential role of metformin in prevention and improve prognosis (Bradley et al., 2018; Higurashi et al., 2016; Hosono et al., 2010; Liu et al., 2017; Ng et al., 2020; Sehdev et al., 2015; Singh et al., 2013). For example, a recent meta-analysis reported metformin use decreased the risk of CRC in T2D patients by 29% and reduced both all-cause mortality of CRC in T2D and CRC-specific mortality in T2D(Wang and Shi, 2021). Furthermore, in *in vitro* models of CRC, metformin has shown synergistic effects with oxaliplatin (Richard and Martinez Marignac, 2015), cisplatin (Zhang et al., 2020), irinotecan (Khader et al., 2021), and fluorouracil (Sang et al., 2020), highlighting metformin as a potential sensitization agent to CRC chemotherapy. Similar observations have been made in *in vitro* models of other cancer types (Berkovic et al., 2021; Fong and To, 2019; Pernicova and Korbonits, 2014).

One possible mechanism of action (MoA) of metformin is the inhibition of mitochondrial respiratory complex I leading to activation of the 5’-AMP-activated protein kinase (AMPK) in a serine/threonine-protein kinase STK11 (LKB1)-dependent manner (Shaw et al., 2005). Thus, AMPK appears to be a widely accepted target of metformin treatment. The activation of AMPK is followed by serine/threonine-protein kinase mTOR (mTOR) inhibition leading to favorable phenotypic outcomes in cancer cells such as reducing protein synthesis and proliferation rates, activation of autophagy, and inhibition of inflammatory responses (Pernicova and Korbonits, 2014). However, AMPK can be also activated by other kinases, such as by the calcium-sensitive kinase calcium/calmodulin-dependent protein kinase kinase 2/beta (CAMKK2) (Hawley et al., 2005; Hurley et al., 2005; Woods et al., 2005) or mitogen-activated protein kinase kinase kinase 7 (MAP3K7; TAK1) during lysosomal injury (Jia et al., 2020). Furthermore, metformin can affect cellular signaling in an AMPK-independent manner (Stein et al., 2019). Therefore, the MoA underlying metformin’s gastrointestinal anti-cancer properties seems pleiotropic and remains largely mysterious (Berkovic *et al*., 2021; Kulkarni *et al*., 2020; Pernicova and Korbonits, 2014; Rena et al., 2017).

The mass spectrometry (MS)-based techniques may facilitate the discovery of MoA for drugs via e.g., phosphoproteomic profiling (Aebersold and Mann, 2016). Regarding the MoA of metformin, a previous phosphoproteomic study in a breast cancer cell line (MCF7) showed that the anti-cancer activity of metformin is not mediated by a limited number of isolated signaling cascades but rather complex (Sacco et al., 2016). In another study, AMPK-dependent and independent kinome perturbation in liver cells treated by metformin uncovered novel kinases mediating hepatic metabolism (Stein *et al*., 2019). As an arising MS technique, data-independent acquisition mass spectrometry (DIA-MS) (Aebersold and Mann, 2016; Gillet et al., 2012; Venable et al., 2004) generates continuous, high-resolution MS2 peak profiles along with liquid chromatography (LC) separation and enables simultaneous identification, localization, and quantification of phosphorylation sites (P-sites) (Bekker-Jensen et al., 2020; Gao et al., 2021; Rosenberger et al., 2017; Wu et al., 2021). Herein, we applied a state-of-the-art DIA-MS-based workflow to perform deep and highly quantitative proteome and phosphoproteome profiling of a panel of 12 highly heterogeneous CRC cell lines treated with metformin. Our study presents a considerable phosphoproteome resource revealing and rationalizing heterogeneity of metformin responses in CRC cells, reinforcing the knowledge of metformin MoA towards cancers therapeutics.

## Results

### A high-quality mass spectrometry dataset captured the acute and late metformin response in CRC

To interrogate the total proteome and phosphoproteome response to metformin in colorectal cancer (CRC) cells, we firstly aimed to select cell lines representing heterogeneous CRC subtypes (**Figure 1A**). Previously, Roumeliotis et al. clustered 50 CRC cell lines based on their proteomic profiles and identified five proteomic subtypes (CPS1-CPS5) which overlapped well with the documented CRC tissue subtypes (Iorio et al., 2016; Medico et al., 2015; Roumeliotis et al., 2017; Sadanandam et al., 2013). In the present study, we therefore selected 12 cell lines to represent these five clusters: three from CPS1 (LoVo, RKO, SW48), five from CPS2 (C2BBe1, HT115, SNU-61, SW948, T84), one from CPS3 (COLO 205), one from CPS4 (MSDT8), and two from CPS5 (NCI-H747, SW837). Next, following a published investigation in MCF7 cells (Sacco *et al*., 2016), we cultured the CRC cells for 24 hours before the treatment with 10 mM metformin (**Figure 1B**) without refreshing the media, to achieve a partial “nutrient exhaustion” state that could be essential to observe AMPK activation (Sacco *et al*., 2016). Unlike Sacco et al., who focused on a 24-hour treatment, we included two time points post metformin addition: 30 minutes for unveiling any potential “acute” phospho-signaling triggered by metformin (Andrzejewski et al., 2014; Ma et al., 2022; Zhang et al., 2012) (measured by phosphoproteome), as well as 24 hours (measured by both proteome and phosphoproteome) for investigating the “late” response of metformin-induced alterations, particularly in cellular metabolism (Sacco *et al*., 2016). We did not measure the proteomic changes at 30 minutes, because such a time window is usually too short for observing any significant protein abundance changes in cancer cells. Moreover, we analyzed time point 0 (i.e., the “steady-state”) with both proteomics and phosphoproteomics as controls. Thus, a total of 108 and 144 samples were randomized for the proteome and phosphoproteome measurements respectively, using a single-shot DIA-MS method (Gao *et al*., 2021; Wu *et al*., 2021).

**Figure 1:**
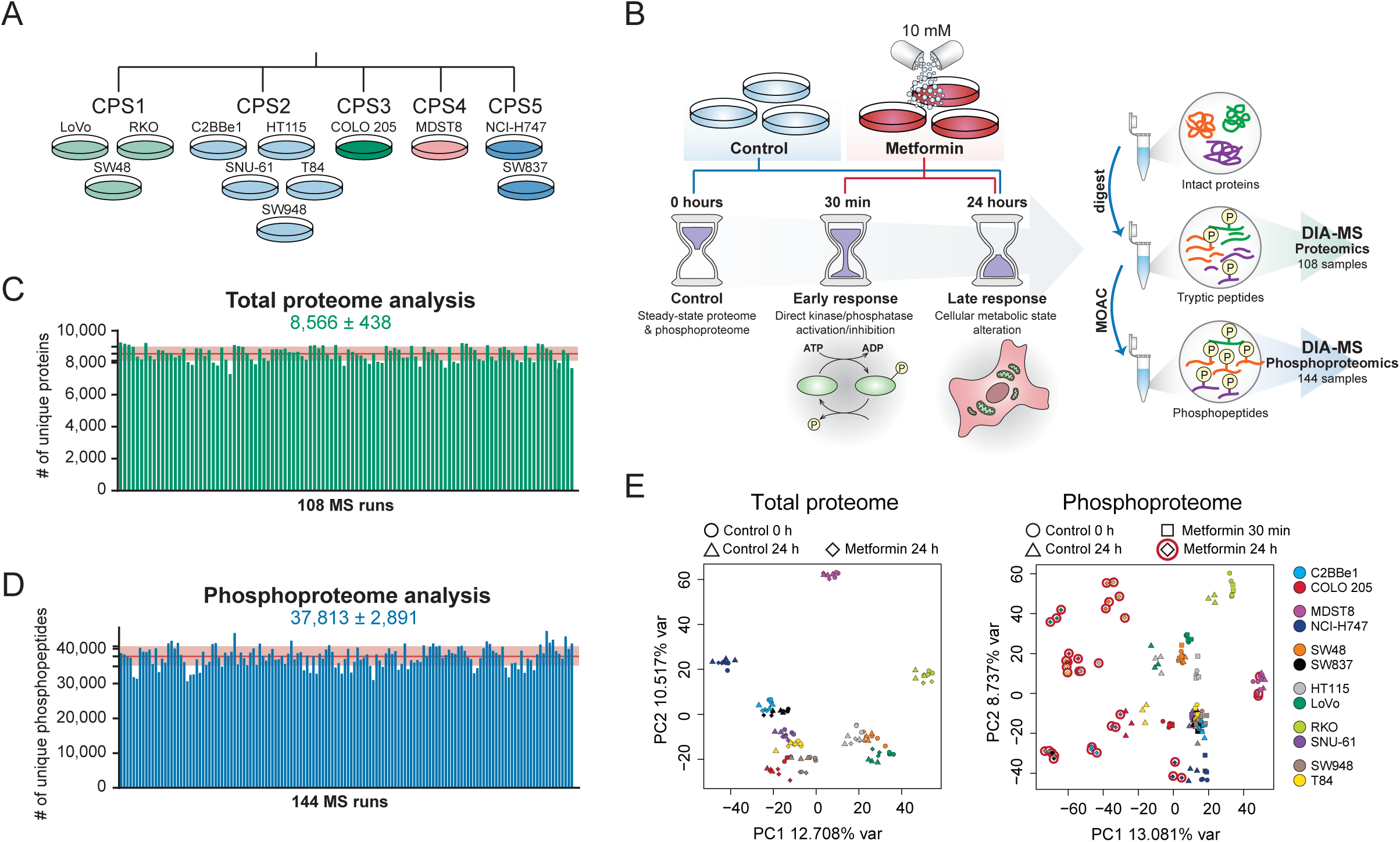
Proteomic and phosphoproteomic analysis of metformin-treated colorectal cancer cells. **(A)** Twelve CRC cell lines selected for the study represent 5 proteomic clusters (CPS1-CPS5) (Roumeliotis *et al*., 2017). **(B)** The cell lines in triplicate dishes per condition were seeded 24 h prior metformin treatment and harvested after 30 min and 24 h to capture the acute and late responses to metformin. Matched tryptic peptides and enriched phosphopeptides were subjected to DIA-MS. Data were analyzed using the library-free algorithm DirectDIA in Spectronaut v14. The PTM workflow in Spectronaut was applied. **(C-D)** The numbers of identified unique protein groups **(C)** and unique phosphopeptides (protein and peptide FDR < 1%) with confidently localized P-sites (i.e., class I sites; **D**) across a total of 252 (i.e., 108+144) randomized injections. **(E)** Principal component analysis of all total proteome (left) and phosphoproteome (right) samples.

Based on a library-free DirectDIA algorithm (Bruderer et al., 2017; Tsou et al., 2015) (see **Methods**), we identified 10,142 protein groups (8,566 ± 438 on average; **Figure 1C**) and 64,680 unique phosphopeptides (37,813 ± 2,891 on average; **Figure 1D**) corresponding to 56,080 unique class I phosphorylation sites (i.e., with localization probability > 0.75; (Bekker-Jensen *et al*., 2020; Olsen et al., 2006)) from 7,450 unique protein groups in the entire experiment, with the peptide- and protein-FDR strictly controlled below 1%. Compared to Sacco et al. in the year of 2016, our phosphoproteomic-DIA (Phos-DIA) profiled 354.6% of their number of P-sites, significantly extending the previous analytical depth (Sacco *et al*., 2016). The median absolute Pearson correlation was 0.93 and 0.88 within replicates for the proteome and phosphoproteome analysis, respectively; and hierarchical clustering analysis of the correlation coefficients showed a predominant clustering based on replicates, indicating a decent reproducibility (**Supplementary Figure 1** & **2**). Moreover, a principal component analysis (**Figure 1E**) confirmed the excellent clustering among biological replicates. To summarize, we acquired deep and highly quantitative DIA-MS datasets capturing the acute and late metformin signaling in 12 representative CRC cells.

### The impact of metformin is much more evident in the phosphoproteome of “late” response

How prevalently metformin affects protein expression? Based on our data, the proteome level change was minor after 24 hours in all CRC cell lines, especially when compared to the phosphoproteome that was extensively rewired at the same time point (**Figure 2A** & **2B**, right panel). Collectively, across all CRC cell lines, while 20.9 ± 12.1% (mean ± SD) of the phosphoproteome was significantly (p < 0.01 & absolute fold change > 1.5) regulated after 24 hours, only 1.7 ± 1.1% of the proteome underwent significant regulation (**Figure 2A** & **2B**, middle panel). In accordance, in PCA the largest variability in the cell line panel at the proteome level was due to the heterogeneity of the baseline proteomes rather than the treatment (**Figure 1E**). In contrast, at the phosphoproteome level, the most variability was explained by different treatment conditions. Interestingly, “control” cells starving for 24 hours demonstrated a notable separation from the time point 0 phosphoproteome samples, confirming the importance of using a time point-matched control. Further correlation analysis (**Figure 2C**) indicated a very weak positive correlation (R = 0.02–0.16) between the proteome and phosphoproteome level. This result suggests the phosphoproteome change was not driven by the total proteome change (which is anyway barely detectable) even for the long-term response. Moreover, the subtraction of the total protein abundance variation from the phosphopeptide variation using linear regression did not show any major impact on the metformin-response quantification (**Supplementary Figure 3**). Therefore, we decided to directly use the phosphoproteomic data, whenever applicable, for the downstream signaling analyses. Altogether, our data suggest metformin only minimally regulates the total proteome expression levels.

**Figure 2:**
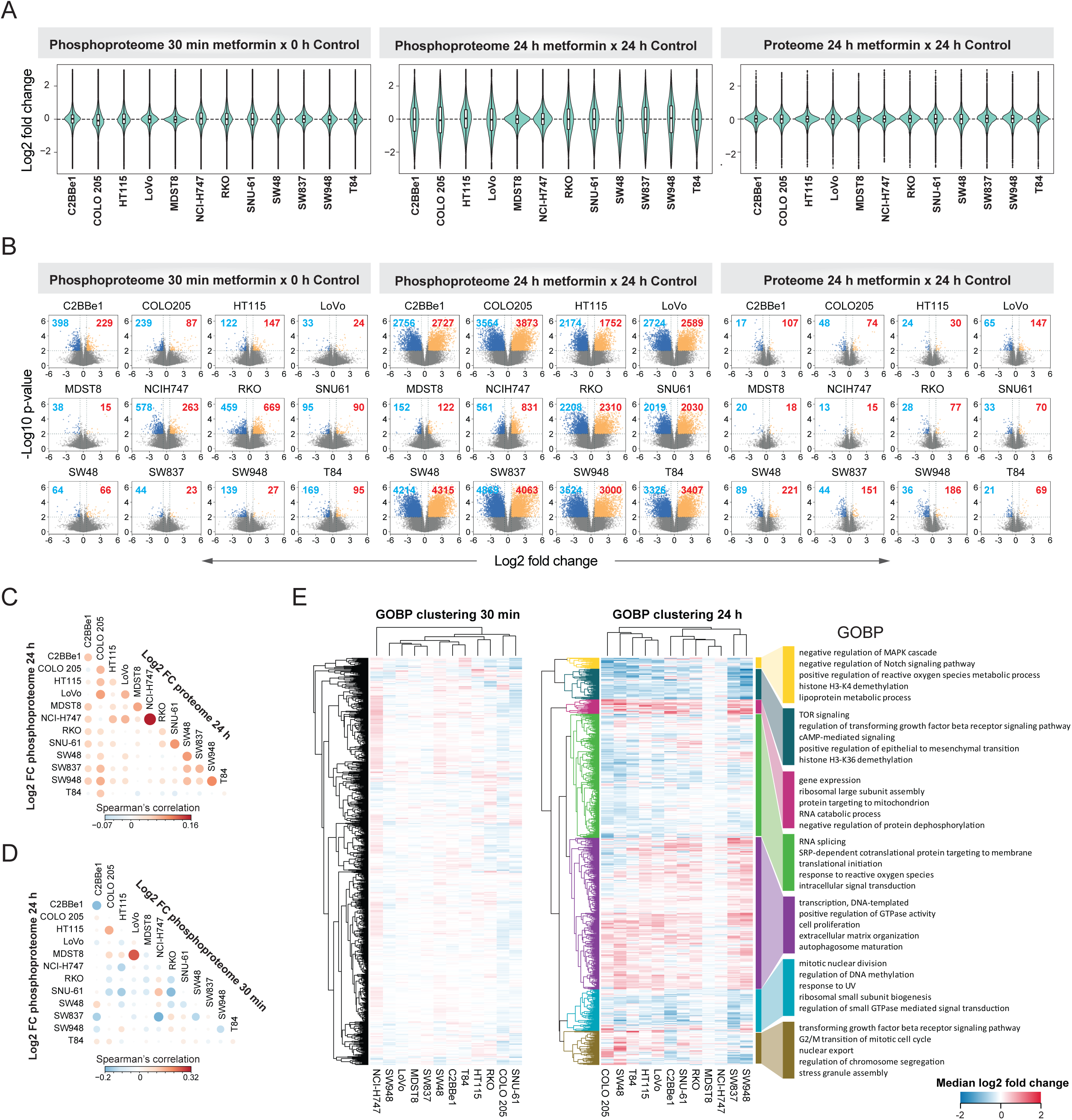
Metformin response occurred predominantly at the phosphoproteome level after 24 hours. **(A)** Violin plots depict the log2 fold change (metformin vs. corresponding control) distribution. **(B)** Volcano plots show the log2 fold change distribution and statistical significance of phosphopeptides and proteins. The blue and red numbers indicate the number of significantly regulated P-sites or proteins per cell line and time point (p < 0.01, absolute fold change > 1.5; two-sided t-test). **(C)** Correlation plot between the 30 min and 24 h log2 fold changes (log2 FC) at the phosphoproteome level. **(D)** Correlation plot between the proteome and phosphoproteome level log2 fold changes after 24 h. **(E)** Hierarchical clustering analysis of gene ontology biological process terms (David GOBP Direct). The processes were prefiltered to at least include 5 unique phosphoprotein ids in each cell line and significantly up- or downregulated in at least one cell line (1D enrichment analysis in Perseus 1.6.14.0). The color illustrates median log2 fold change per GOBP in each cell line based on all phosphoproteins in annotated by a GOBP. The same set of GOBPs in the 30 min (left) and 24 h (right) heatmaps are shown. The right panel shows selected significant GOBP terms per GOBP cluster based on the 24 h data.

Previous studies have suggested the existence of an acute response to metformin treatment (Andrzejewski *et al*., 2014; Ma *et al*., 2022; Zhang *et al*., 2012). However, phosphoproteomic profiling of the rapid response has been lacking. As shown in **Figure 2A** and **2B** (left panel), we identified differential P-sites following the 30-minute treatment underscoring an “acute” response of a fraction of the phosphoproteome. However, both the magnitude of the change and the number of significant phosphopeptides were markedly lower compared to the “late” response, after 24 hours. Specifically, we found the most prominent acute phosphorylation response in RKO, NCI-H747, and C2BBe1. Interestingly, for any of these cell lines, the acute and late response did not correlate (**Figure 2D**) and affected a distinct set of phosphorylation sites. The low extent of phosphoproteome perturbation after 30 minutes was also reflected at the level of signaling pathways modulated by metformin treatment (**Figure 2E**, left panel). These results are suggestive of distinctive temporal patterns in the rapid metformin response of particular CRC cells.

In summary, the dominant metformin-induced response occurred at the phosphoproteome level rather than at the proteome expression level in CRC, and a long-term 24-hour treatment showed a more extensive perturbation of the phosphoproteome.

### The P-site response to metformin is highly heterogenous among CRC cells

The 12 cell lines were selected to represent variable CRC genotypes and highly heterogenous signaling repertoires (Roumeliotis *et al*., 2017). We thus interrogated how the cell line variability affects the metformin response after 24 hours of the treatment. ***Firstly***, the fold change distributions (**Figure 2A**) and volcano plots (**Figure 2B**) clearly distinguished between two groups of cells – in most (10 of 12) cell lines, we identified thousands of differentially phosphorylated peptides (∼13.8-41.9% of the phosphoproteome measured per cell line), while in the other two cell lines, MDST8 and NCI-H747, we discovered only minor phosphorylation change (0.8% and 4.1%). Furthermore, even for the former 10 cell lines, the overlaps were rather minor (0.08% of the measured phosphoproteome, **Supplementary Figure 4**). ***Secondly***, to further classify the cell lines, we performed a consensus clustering analysis using the top 30% most variable phosphopeptides based on metformin induced log2 fold change across cell lines. We identified 4 consensus clusters (**Supplementary Figure 5A**), which interestingly did not resemble the clusters identified in the steady-state proteome and phosphoproteome data (**Supplementary Figure 5B**), suggesting that the heterogenous metformin response is not easily predictable from the basal state. ***Thirdly***, we sought to investigate the heterogeneity of the biological processes induced by metformin between cell lines. Based on the 1D enrichment analysis (**Figure 2E**), we found that processes such as gene expression, RNA splicing, and translational initiation were enriched in upregulated direction in most of the cell lines (in all cell lines 1D enrichment *p*-values < 0.05), indicating that CRC cells prefer to trigger these pathways to cope with the metformin-altered cellular metabolism. On the other hand, among the processes with downregulated phosphorylation, TOR signaling, mitotic nuclear division, or regulation of DNA methylation were processes significantly enriched in most of the cell lines. However, at least half of the processes did not follow the same trend among the 12 cell lines (**Figure 2E**).

In conclusion, our data suggests that while most cell lines altered the phosphoproteomes as a result of drug perturbation, the degree of the alteration, the specific phosphorylation sites, and the biological processes triggered or inhibited by metformin were highly variable across the CRC panel.

### The lysosomal impact of metformin according to quantitative proteomics and phosphoproteomics

A recent study identified key lysosomal metformin-protein interactors, elucidating the mechanism of the lysosomal glucose-sensing pathway leading to AMPK activation (Ma *et al*., 2022). Although we did not measure the metformin-protein interactions directly, we mapped our quantitative proteome and phosphoproteome data to the metformin interacting lysosomal protein list in Ma et al. (n = 87, hereafter, “Ma list”; see **Methods**). Intriguingly, while the protein abundances of Ma list identities did not change post metformin treatment (**Figure 3A**), in 9 out of 12 cells we found the upregulation of the P-sites (n = 114) for the Ma list proteins (*p* = 0.0495 to 3.1e-06) upon the 24-hour drug treatment (**Figure 3B**). This result thus implicates that metformin-interacting proteins might have increased phosphorylation levels, propagating to subsequent signaling transduction.

**Figure 3:**
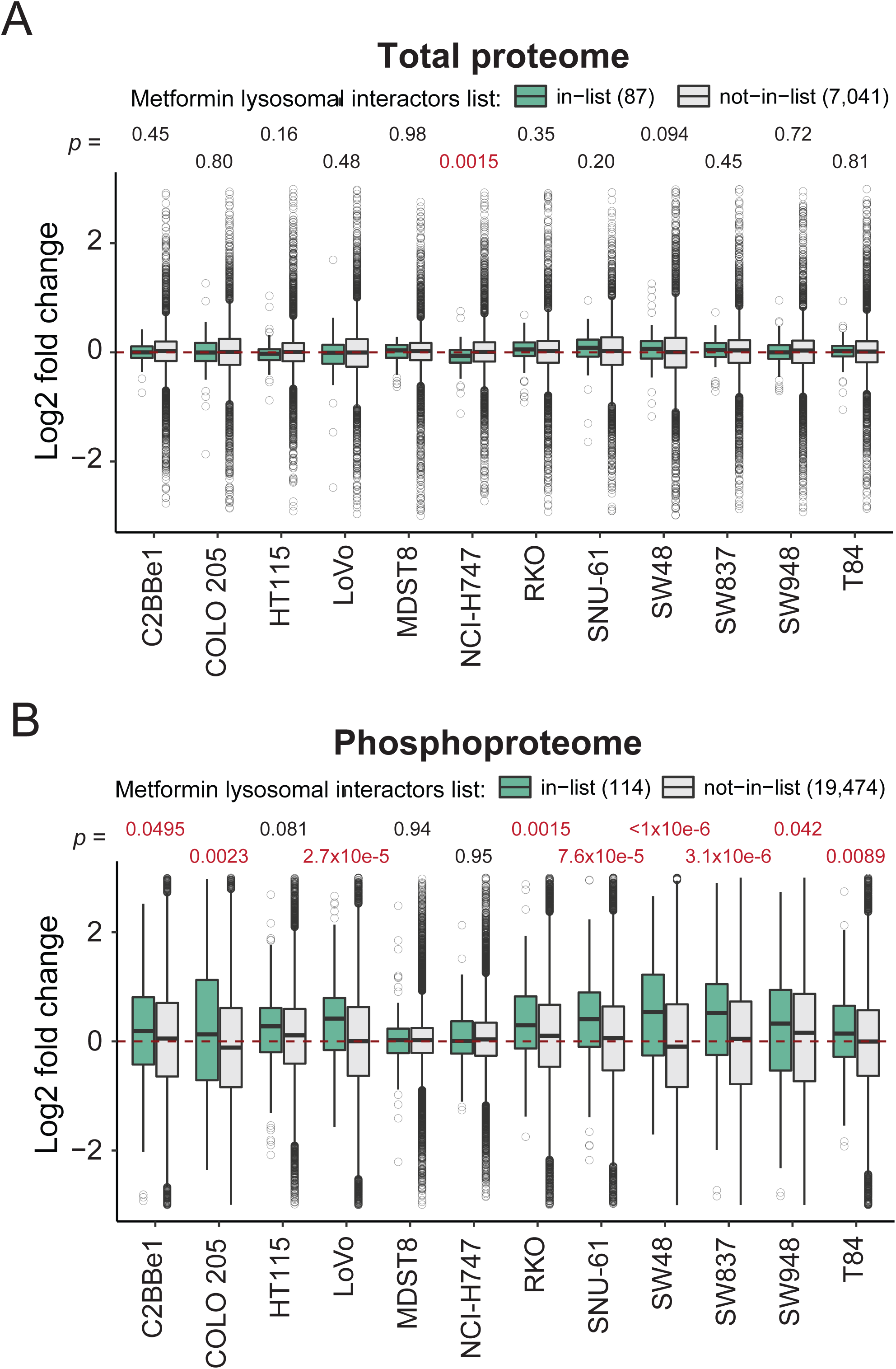
Metformin-interacting lysosomal proteins tend to be increasingly phosphorylated after 24 hours. **(A)** The total proteome and the **(B)** phosphoproteome log2 fold changes induced by metformin treatment. The list of metformin-interacting proteins (green boxes) is based on a recently published dataset (Ma *et al*., 2022). Statistical analysis was performed using the Wilcox test.

### Establishing MetScore, a score summarizing metformin association for each P-site

Based on the heterogeneity of the metformin response among 12 cells, we contemplated that assigning a consensus score to each P-site quantifying its responsivity across the panel would enable the functional stratification of the P-sites. Additionally, P-sites perturbed by metformin in all or most of the cell lines could be curated as a CRC-specific “metformin signature”. To do so, we devised a scoring strategy named “MetScore” (**Figure 4A**). For every P-site, a “MetScore” is assigned as a sum of cell lines in which a P-site was significantly upregulated (+1), non-regulated (+0), or downregulated (-1). MetScore stratified all P-sites into 5 segments, G1 to G5, with G1 encompassing the mostly upregulated P-sites (in at least 4 CRC cell lines) and G5 encompassing P sites downregulated in at least 4 lines. Intriguingly, the P-sites in the G1, G2, and G5 MetScore segments showed on average significantly larger *site-specific functional scores* (Ochoa et al., 2020) than the P-sites in the G3 segment. This result supports the greater functional relevance of P-sites with extreme MetScores, as compared to the unperturbed sites **(Figure 4B**). Remarkably, a small list of P-sites (n = 43 and 12) were consistently up- or downregulated in at least 10 cell lines (absolute MetScore ≥ 10; **Figure 4A**), respectively (**Table 1**). The only one P-site with a positive MetScore of 12 (upregulated in all cell lines) was the Progesterone receptor membrane component 2 (PGRMC2) Ser 104. According to the PhosphoSitePlus database (Hornbeck et al., 2012; Hornbeck et al., 2015), the Ser 104 P-site has been identified in multiple high-throughput studies; however, it is not known whether the P-site carries any regulatory function. Strikingly, the PGRMC2 protein is a trans-membrane progesterone receptor that has been recently proposed as an essential player in adipocyte glucose metabolism (Galmozzi et al., 2019). Additionally, two P-sites with MetScore = 11, Armadillo repeat-containing protein 10 (ARMC10) Ser 45, and Stromal interaction molecule 1 (STIM1) Ser 257 have been respectively validated as modulators for AMPK mediated mitochondrial dynamics (Chen et al., 2019) and for intracellular calcium flux during exercise (Nelson et al., 2019). On the other side, the Period circadian protein homolog 2 (PER2) Ser 977 P-site was one of the most commonly downregulated sites (MetScore = -11). The potential function of the P-site has not been described yet; however, the circadian regulator PER2 has been previously shown to also exert important functions in controlling lipid metabolism (Grimaldi et al., 2010). Functional analyses for P-sites with highest and lowest MetScores are therefore warranted in the future to understand their associations with metformin.

**Figure 4:**
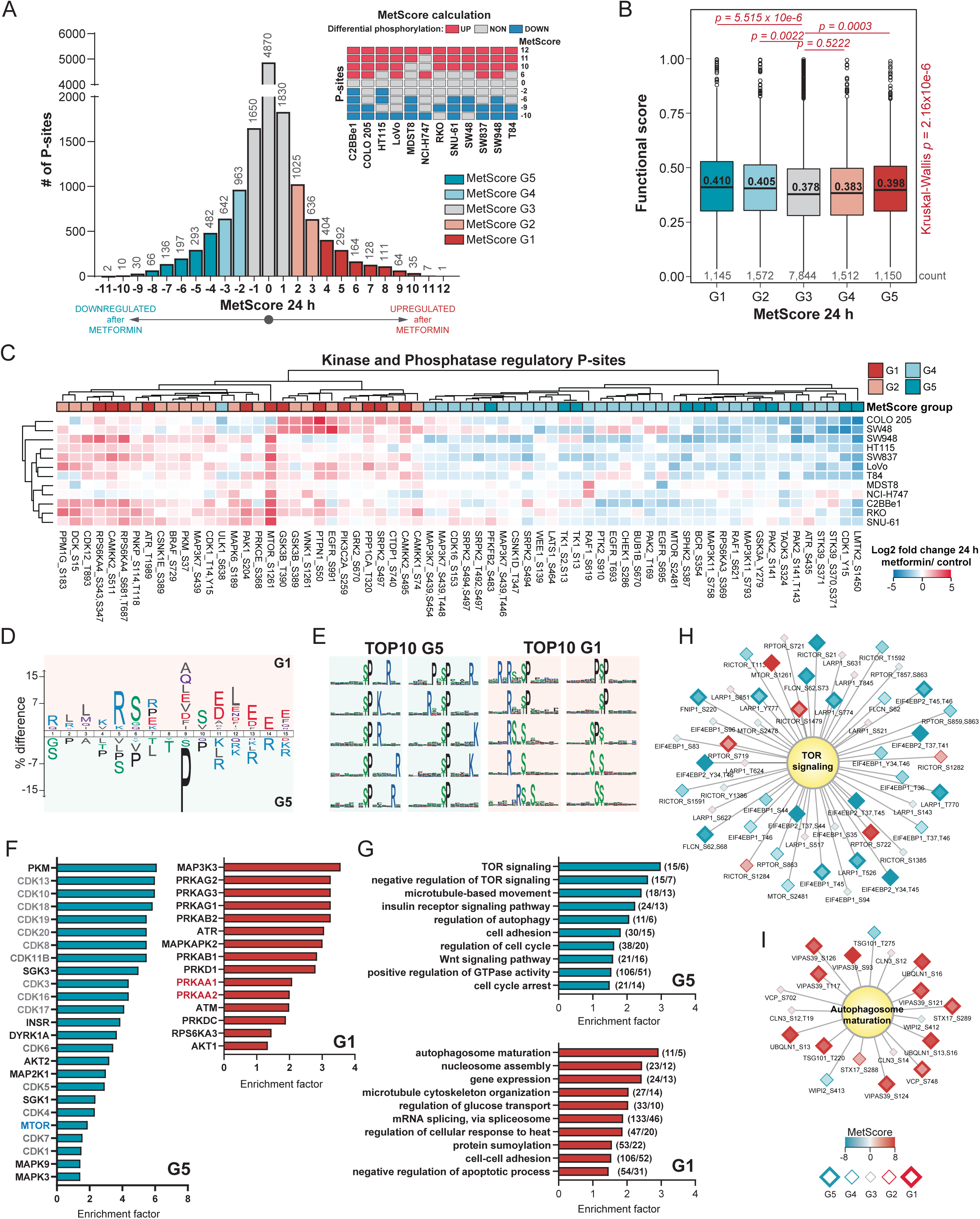
Identification and functional characterization of the common metformin phosphorylation signature using MetScore. **(A)** MetScore was calculated for each P-site based on the results of the statistical analysis (two-sample t-test *p* < 0.01 & absolute fold change > 1.5) as a sum of cell lines in which a P-site was upregulated (+1), non-regulated (+0), or downregulated (-1). P-sites with missing statistical analysis results (i.e., with less than 2 valid values per group) were excluded. Histogram shows the MetScore distribution of all P-sites (n = 14,032). The P-sites were divided into 5 segments for a subsequent functional analysis. **(B)** Distribution of the *P-site specific functional scores* based on a previous publication by Ochoa et al. (Ochoa *et al*., 2020) in the MetScore segments. Statistical analysis was performed using Kruskal-Wallis test; the pairwise comparisons were performed using pairwise Wilcox test with Benjamini-Hochberg correction. **(C)** Hierarchical clustering analysis of the “regulatory” P-sites (based on PhosphoSitePlus database) from kinases and phosphatases with an absolute MetScore ≥ 2. **(D)** Sequence analysis was performed using iceLogo comparing the relative frequencies of amino acids at different positions in the sequence window between the G1 group (foreground/top) and G5 group (background/bottom, *p* < 0.05 as estimated by iceLogo). The size of the of an amino acid in the figure reflects the difference in the frequency between G1 and G5. **(E)** Top10 most significantly enriched sequence motifs extracted using motifeR (minimal occurrence = 20 & *p* < 0.000001) from the 15 A.A. sequence windows surrounding the P-sites in G1 and G5. **(F)** Selected most significant results of kinase-substrate enrichment analysis (Fisher’s exact test *p* < 0.05). The site-specific kinase-substrate annotation was retrieved from OmniPath database. **(G)** Selected most significant results of the protein-level annotation gene ontology biological process (GOBP) terms enrichment analysis (Fisher’s exact test *p* < 0.05; relative to the number of phosphoproteins per category). Number of phosphorylation sites and phosphoproteins are shown in brackets. **(H-I)** Networks of P-sites corresponding to the TOP most significant GOBP terms from **(G)** are depicted. The node fill color indicates the MetScore; the border and node size indicate the MetScore segment.

**Table 1:** P-sites identified as the common metformin signature. Selected P-sites with absolute MetScore ≥ 10. PSPdb “+” indicates the site is listed in the PhosphoSitePlus database (January 2022). Reg. site “+” indicates the site has a known regulatory function in the database.

| MetScore | Gene name | P-site | PSPdb | Reg.<br>site | MetScore | Gene name | P-site | PSPdb | Reg.<br>site |
| --- | --- | --- | --- | --- | --- | --- | --- | --- | --- |
| 12 | PGRMC2 | S104 | + |  | 10 | RANBP2 | S1955 | + |  |
| 11 | ARMC10 | S45 | + | + | 10 | SAFB2 | T193 |  |  |
| 11 | ASPSCR1 | S502 | + |  | 10 | SCAMP2 | S319 | + |  |
| 11 | BCLAF1 | S196 | + |  | 10 | SCAMP2 | S320 | + |  |
| 11 | GPAT3 | S68 | + |  | 10 | SCRIB | S1276 | + |  |
| 11 | PRPF4B | S8 | + |  | 10 | SLC12A2 | S77 | + | + |
| 11 | SRRM2 | T367 | + |  | 10 | SNX27 | S51 | + |  |
| 11 | STIM1 | S257 | + | + | 10 | SRRM2 | T359,T367 | + |  |
| 10 | ANXA2 | S2 | + |  | 10 | SRSF6 | S119 | + |  |
| 10 | ANXA2 | T3 | + |  | 10 | SUGP1 | S409 | + |  |
| 10 | ARHGEF11 | S663 | + |  | 10 | TNKS1BP1 | S1620 | + |  |
| 10 | BAD | S99 | + | + | 10 | UGDH | S476 | + |  |
| 10 | CFL1 | S8 | + |  | 10 | ZC3H13 | S986 | + |  |
| 10 | CLINT1 | S166 | + |  | 10 | ZC3H13 | S370,S372 | + |  |
| 10 | EEA1 | S52 | + |  | 10 | ZC3H13 | S370,Y374 | + |  |
| 10 | EEF2 | S502 | + |  | -10 | BAHD1 | S184 | + |  |
| 10 | EIF5 | S10 | + |  | -10 | JPT2 | S97 | + |  |
| 10 | FIP1L1 | S492,T494 | + |  | -10 | LASP1 | T104 | + |  |
| 10 | HMGA1 | S49 | + |  | -10 | LMTK2 | S1450 | + | + |
| 10 | HNRNPH1 | S23 | + |  | -10 | MTDH | S568 | + |  |
| 10 | HNRNPM | S432 | + |  | -10 | RBMX | S208 | + |  |
| 10 | MAP4 | S928 | + | + | -10 | RPRD2 | S758,T763 | + |  |
| 10 | MAP7D1 | S313 | + |  | -10 | SF3B1 | T426 | + |  |
| 10 | NUFIP2 | S652 | + |  | -10 | STMN1 | S38 | + | + |
| 10 | PCYT1A | S331 | + |  | -10 | TLE3 | T334 | + |  |
| 10 | PHRF1 | S814 | + |  | -11 | PER2 | S977 | + |  |
| 10 | PTMA | S10 | + |  | -11 | ZDHHC5 | S621 | + |  |
| 10 | RANBP1 | S188 | + |  |  |  |  |  |  |

Following, we assessed known regulatory P-sites in kinases and phosphatases using their MetScores in more detail (**Figure 4C**). A total of 66 regulatory kinases or phosphatase P-sites were identified and significantly enriched in G1, G2, G4, and G5 segments compared to those unperturbed P-sites in the G3 segment (Fisher’s exact test *p*-value = 0.0072). Among them, 36 were annotated as regulating enzymatic activity by PhosphoSitePlus. In particular, we found two mTOR regulatory sites, Ser 1261 and Ser 2841. While the AMPK substrate mTOR Ser 1261 (Acosta-Jaquez et al., 2009) was mostly upregulated with MetScore = 8, the “classic” mTOR-activating autophosphorylation site, Ser 2841, had a MetScore of -2. Thus, Ser 2481 appeared to be not significantly modulated by metformin in most CRC cell lines – this observation agrees well with the result in MCF7 cells, and therefore reinforces the conclusion of Sacco et al. that PI3K-mTOR pathway was rewired rather than completely turned off upon 24-hour metformin treatment in cancer cells (Sacco *et al*., 2016).

Next, we performed a series of functional enrichment analyses for G1 and G5 segments. ***Firstly,*** we compared the amino acid sequences surrounding the G1 and G5 sites using iceLogo (p < 0.05; **Figure 4D**) (Colaert et al., 2009) revealing that basic amino acids (Arg and Lys) at the positions preceding the P-sites and acidic amino acids (Glu and Asp) following the P-sites were enriched in the G1 segment. On the other hand, there was a relative enrichment of proline at +1 position and basic amino acids (Arg and Lys) at the positions following a P-site in G5. The strong preference for proline-directed motifs in G5 indicated that metformin preferentially downregulated proline-directed protein kinases such as mitogen-activated protein kinases (MAPKs) and cyclin-dependent kinases (CDKs). Reassuringly, the motif enrichment results by motifeR (**Figure 4E**) (Wang et al., 2019) showed that all top 10 most significant motifs in G5 have a proline at +1 position (**Figure 4E**), including the S/T-P-x-K/R, the classic substrate motif for CDKs (Songyang et al., 1994), as well as P-x-S/T-P, the established motif for MAPKs (Gonzalez et al., 1991). In contrast, an AKT substrate motif, R-x-R-x-x-S/T (Alessi et al., 1996), was enriched in G1 sites. ***Secondly*,** we performed an enrichment analysis for known kinase targets extracted from the OmniPath database (Ceccarelli et al., 2020; Türei et al., 2016; Türei et al., 2021) (Fisher’s exact test *p* < 0.05; **Figure 4F**). In addition to CDKs enriched in G5, this analysis revealed mTOR substrates were enriched in G5, while the AMPK (PRKAA1) substrates were enriched in G1, confirming the importance of AMPK-mTOR axis following metformin treatment. The G1 segment also enriched substrates of kinases previously reported to be activated by metformin in the previous two phosphoproteomic reports (Sacco *et al*., 2016; Stein *et al*., 2019), such as AKT Serine/Threonine Kinase 1 (AKT1), MAPK Activated Protein Kinase 2 (MAPKAPK2; MK2), or protein kinase D (PRDKD1; PKD). ***Thirdly,*** we analyzed significant GO BPs in G1 and G5 segments at the P-site level. We identified terms related to mTOR signaling, autophagy regulation, cell proliferation, and general cellular processes such as gene expression, mRNA splicing, and protein modifications, many are overlapping to **Figure 2E** (i.e., the protein-level enrichment), as expected (Fisher’s exact test p < 0.05; **Figure 4G**). As particular examples, most P-sites in proteins involved in mTOR signaling were downregulated whereas the autophagosome maturation process contained predominantly upregulated sites (**Figure 4H-I**). We additionally identified a significant overrepresentation of the Wnt signaling pathway in G5, which has been shown to be hyperactivated in almost all CRCs as an initiating event (Zhan et al., 2017). Hypophosphorylation in the Wnt pathway upon metformin might indicate a potential anti-CRC mechanism of metformin.

To summarize, our multi-cell line data in CRC were used to assign a unique MetScore per each P-site in the phosphoproteome, which facilitated the P-site specific functional analysis towards the in-depth understanding of metformin signaling.

### Inferring the kinase activity landscape and signaling nodes in CRC cells following metformin treatment

To infer the kinase activity response in a cell line-specific manner, we performed an enrichment on the phosphoproteomic data using decoupleR (Badia-i-Mompel et al., 2022), leveraging the kinase-P-site interactions from the OmniPath database (Türei *et al*., 2016; Türei *et al*., 2021). The group of kinases that showed mostly downregulated activities across the cell lines (**Figure 5A**, green cluster) include many kinases regulating cellular growth and proliferation, such as mTOR, CDKs, mitotic regulators AURKA, AURKB, and PLK1, MAPKs, but also several tyrosine kinases such as EGFR, FYN, and ABL1 and a dual specificity kinase DYRK1A, which is not evident from the above P-site level analyses. On the other hand, the cluster of kinases with a stronger upregulation in the consensus cluster 1 cell lines (i.e., RKO, SNU-61, HT115, SW948, C2BBe1, SW48; **Figure 5A**, purple cluster), but not in all cells, harbors AMPK (PRKAA1) as well as other metabolic regulators such as the insulin receptor (INSR), a tyrosine kinase regulating glucose metabolism. Remarkably, this cell consensus cluster also showed increase activities for the upstream CDKs regulators in response to different stress stimuli such as ATM, ATR, the catalytic subunit of DNA-PK (PRKDC), and the downstream effector of ATR, CHEK1, potentially explaining a stronger downregulation of CDKs in these cell lines. Serine/threonine-protein kinase STK11 (STK11; LKB1), a protein kinase essential for AMPK activation under low ATP conditions (Shaw *et al*., 2005) was determined to be activated in most of the cell lines (**Figure 5A**, orange cluster). The cell line-specific, metformin-induced kinase activity landscape is further illustrated by the kinome trees (**Figure 5B**). The color mapping of branches and nodes corresponding to the kinase activity scores highlight variable fingerprints of cell signaling states in e.g., the CAMK and AGC kinome branches. Surprisingly, in the COLO 205 cell line, the reduced activity was inferred for most kinases, while there does not seem to be any preference for P-site downregulation in COLO 205 (see the volcano plot in **Figure 2B**). This discrepancy supports the value of kinase activity inference to analyze the phosphoproteomic data.

**Figure 5:**
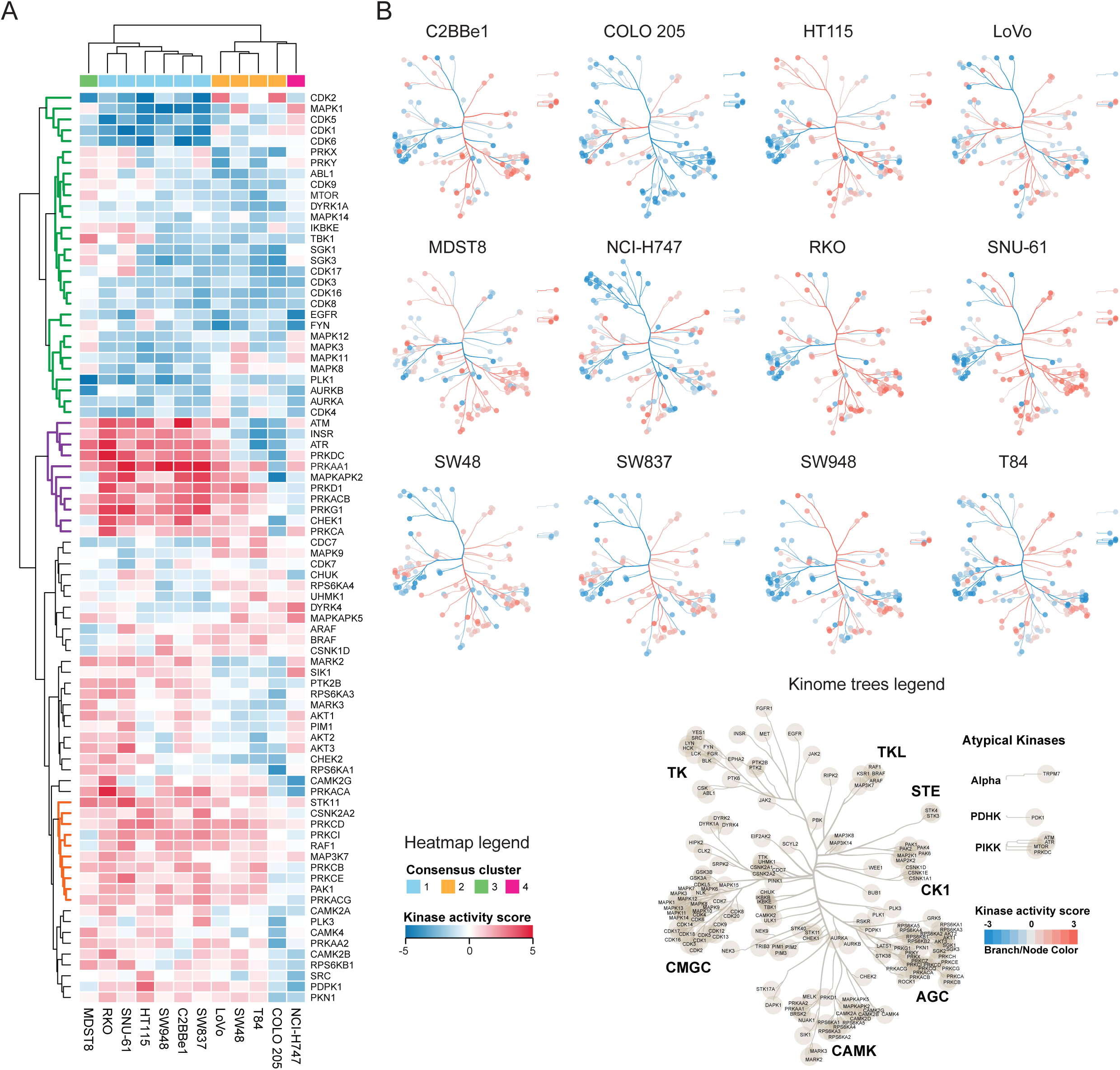
Kinase activity landscape highlights the heterogeneity of metformin response across the cell lines. **(A)** Hierarchical clustering analysis of kinase activity scores estimated using decoupleR rewired after 24 h of metformin treatment. Only kinases with statistically significant activity scores (*p* < 0.05) in at least one cell line and estimated activities in all 12 cell lines are shown. **(B)** The metformin-rewired kinomes were visualized per cell line. The branch and node color mapping correspond to the kinase activity score. Only those nodes and branches with a valid score per cell line are shown. The legend shows all branches and kinases with computed activity scores in at least one cell line.

To derive mechanistic insights into MoA of metformin in individual cell lines, we contextualized the networks using PHONEMeS (Gjerga et al., 2021; Terfve et al., 2015). Three inputs were used for the analysis (**Figure 6A**), i) the top 15% of differentially expressed P-sites, ii) the top 15% of differentially activated kinases, and iii) the prior knowledge network extracted from OmniPath database consisting of protein-protein and kinase-P-site interactions (Türei *et al*., 2016; Türei *et al*., 2021). Using these inputs, PHONEMeS reconstructed a coherent path connecting deregulated kinases with deregulated P-sites resulting in large signaling networks (**Supplementary Figure 6A-L**). These networks contain the perturbed phosphoproteome and kinases (from the inputs), as well as the inferred intermediate nodes and a computed score reflecting the activity of each node (**Figure 6A**). As an example of prominent modules in these networks, we focused the subnetwork on AMPK and its closest upstream regulators and downstream effectors to compare the signaling pathways leading to AMPK regulation across cell lines. The reconstructed subnetworks illustrated the substantial differences between the cell lines in their response to metformin after 24 hours (**Figure 6C)**. ***First,*** AMPK, as the central node, was not perturbed the same in all the cell lines. In MDST8, a cell line mostly non-responding at the phosphoproteome level, AMPK was not activated. In COLO 205, the outlier in the kinome visualization (**Figure 5B**), AMPK activity was inferred to be downregulated. The rest of the cell lines showed AMPK activation, albeit with a different extent. ***Second*,** the upstream signals leading to AMPK activation seem to be divergent between cells. Previously, AMPK activity was reported to be regulated by several kinases in a context-dependent manner (Steinberg and Carling, 2019). For example, in the context of low energy (ATP) conditions, STK11 is essential for AMPK phosphorylation at Thr 172 and AMPK activation (Shaw *et al*., 2005). Under conditions of calcium flux, however, despite the same residue Thr 172 is phosphorylated, AMPK activity is regulated by the calcium-sensitive kinase calcium/calmodulin-dependent protein kinase kinase 2/beta (CAMKK2) (Hawley *et al*., 2005; Hurley *et al*., 2005; Woods *et al*., 2005). Furthermore, mitogen-activated protein kinase kinase kinase 7 (MAP3K7; TAK1) controls AMPK during lysosomal injury by various agents including metformin treatment (Jia *et al*., 2020). Our reconstructed signaling networks (**Figure 6C**) suggested STK11 was upstream of AMPK in three cell lines, while the CAMKK1 and CAMKK2 kinases were acting upstream in nine cell lines. Meanwhile, in five cell lines the MAP3K7 was found to be the upstream AMPK regulator. These results largely support the context-dependent AMPK activation. ***Third,*** the activity of the downstream AMPK nodes also showed a considerable heterogeneity. Half of the cell lines showed mTOR inactivation followed by inhibition of autophagy inhibitor serine/threonine-protein kinase Sgk1 (SGK1) (Hart et al., 2016) as the best path explaining the signaling outcome in four cell lines (**Figure 6C**). Other downstream effectors were usually not shared by more than one or two cell lines.

**Figure 6:**
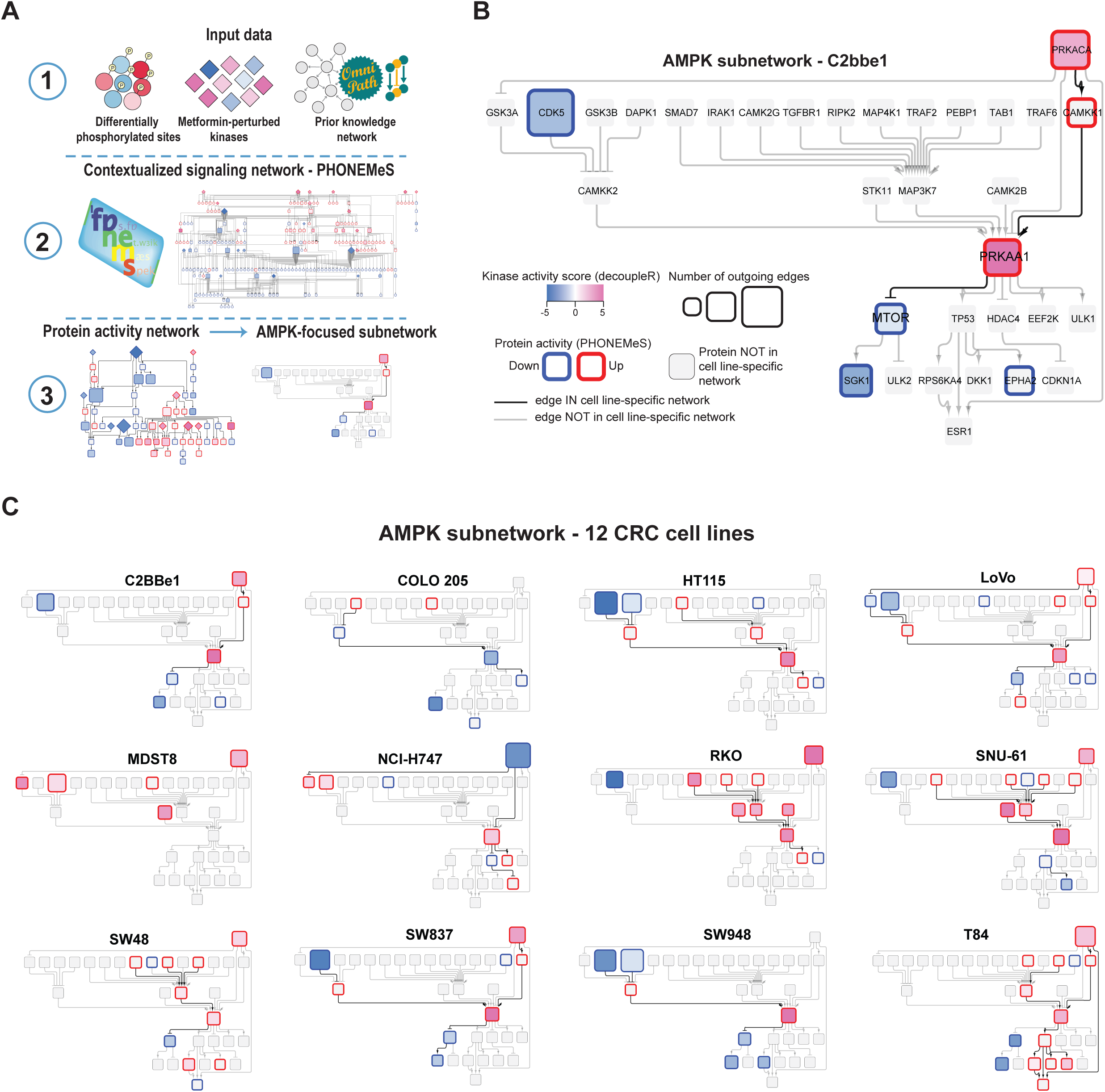
Network analysis using PHONEMeS reveals a substantial heterogeneity in the AMPK-focused subnetwork across CRC cells. **(A)** PHONEMeS analysis workflow. The results of the differential analysis (t-value; limma), the results of the kinase activity analysis (normalized kinase activity score, decoupleR), and prior knowledge network containing protein-protein and kinase-P-sites interactions retrieved from OmniPath were used as an input for PHONEMEeS to reconstruct cell line-specific signaling networks. A general AMPK-focused subnetwork was constructed using all 2-step up- and down-stream neighbors of AMPK included in the network in at least one of the cell lines. The cell line-specific nodes and edges were highlighted. **(B)** An example of C2BBe1 AMPK-focused subnetwork. The border color corresponds to the protein activity scores calculated using PHONEMeS; the fill color mapping illustrated the decoupleR kinase activity score. The bold edges are part of the C2BBe1 subnetwork. **(C)** Subnetworks from all 12 CRC cell lines illustrate the cell-specificity of metformin-induced signaling. The position of the nodes (proteins) and edges is the same as in **(B).**

Altogether, using phosphoproteomics-tailored bioinformatic tools, the metformin-signaling leading to AMPK activation and downstream signal propagation were revealed to be orchestrated by different kinase activities and dependent on specific cellular context.

### Leveraging the metformin signature to predict novel metformin-drug interactions

Recently, several computational tools and resources have enabled annotating specific P-site as a known drug target (Krug et al., 2019) or correlating the P-site abundance (Abelin et al., 2016; Dele-Oni et al., 2021; Litichevskiy et al., 2018) with drug sensitivity across cell line panels (Frejno et al., 2020). Furthermore, metformin has shown synergistic effects with CRC genotoxic chemotherapy *in vitro (*Khader *et al*., 2021; Liu et al., 2020; Richard and Martinez Marignac, 2015; Sang *et al*., 2020). Therefore, we asked whether we could leverage the metformin phosphoproteomic signatures we discovered to predict the metformin-drug interactions in CRC. To do so, we focused on the 55 P-sites with an absolute MetScore ≥ 10 (**Figure 4A** & **Table 1**) and queried them separately in the ATLANTiC website (Frejno *et al*., 2020), which hosts significant drug-sites association predicted based on the CRC panel of 65 cell lines (from here on, the CRC65 dataset). From CRC-specific drug-sites associations, we extracted nine compounds with different targets and mechanism of action significantly associated with the most signature P-sites (**Figure 7**), i.e., cetuximab (EGFR inhibitor), paclitaxel (mitosis inhibitor), pazopanib (tyrosine kinase inhibitor), navitoclax (Bcl-2 family protein inhibitor), nilotinib (Bcr-Abl tyrosine kinase inhibitor), nutlin-3 (p53-MDM2 inhibitor), SN-38 (active metabolite of topoisomerase I inhibitor irinotecan), vorinostat (HDAC inhibitor), and YM-155 (survivin inhibitor and autophagy activator). Among them, cetuximab, paclitaxel, pazopanib, nilotinib, and vorinostat are FDA approved cancer drugs for various indications, including metastatic CRC (in the case of cetuximab). Additionally, navitoclax, SN-38, and YM-155 are promising compounds being evaluated in clinical trials. Favorably, several drugs in this list have been shown to interact with metformin in *in vitro* models of several cancer types. For instance, as for paclitaxel, metformin has been reported to sensitize ovarian, endometrial, melanoma, and prostate cancer cells to paclitaxel treatment (Dos Santos Guimaraes et al., 2018; Hanna et al., 2012; Ko et al., 2019; Lee and Park, 2021; Lengyel et al., 2015; Zhao et al., 2019). As for nilotinib, metformin has increased the efficacy of nilotinib in chronic myelogenous leukemia cell lines (Na et al., 2021). As for vorinostat, metformin and vorinostat have shown a synergy in overcoming EGFR tyrosine kinase inhibitor resistance in non-small cell lung cancer (Chen et al., 2017). And finally, as for pazopanib, in a clinical study, the use of metformin was associated with favorable outcome of metastatic renal cell carcinoma patients treated with pazopanib (Fiala et al., 2021). In addition to ATLANTiC associations, the HDAC inhibitor vorinostat was enriched in P-site annotations using the PTMsigDB (Krug *et al*., 2019), whereas both vorinostat and paclitaxel emerged when correlating the individual cell line-specific metformin signatures to the drug profiles using the P100 (Abelin *et al*., 2016; Dele-Oni *et al*., 2021; Litichevskiy *et al*., 2018) and PhosFate Profiler (Ochoa et al., 2016) (**Supplementary Figure 7**).

**Figure 7:**
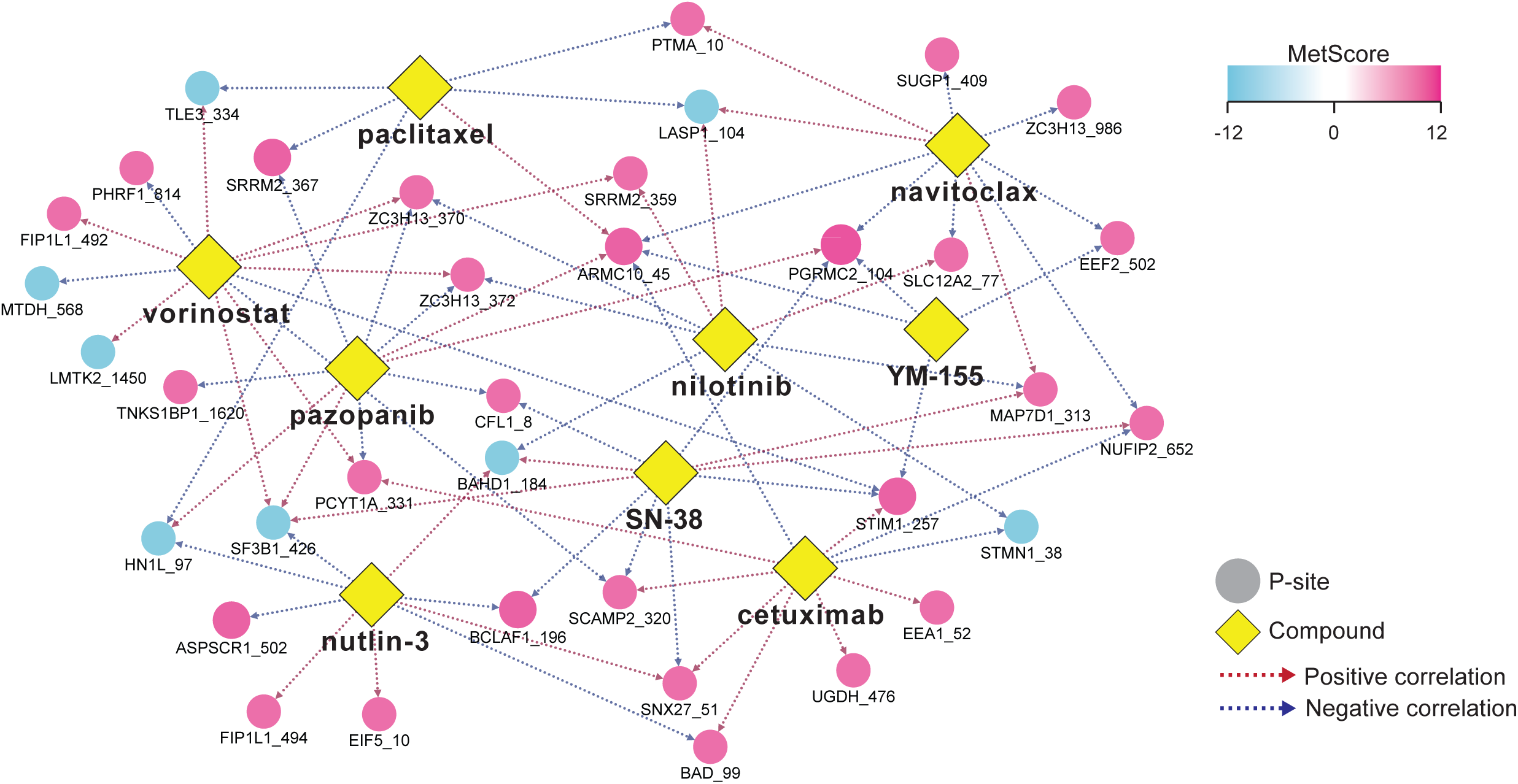
Identification of CRC-specific metformin-drug interactions. **(A)** P-site∼drug network was reconstructed using significant associations of the TOP and BOTTOM MetScore P-sites (absolute MetScore ≥10) retrieved from ATLANTiC database. The drugs/compounds are highlighted by yellow diamonds; the P-sites are shown as circles. The color of the circles indicates the MetScore. The color of the edges indicates whether the abundance of a P-site was shown to be significantly positively or negatively correlated with a sensitivity to a drug in the ATLANTiC database.

In conclusion, leveraging our MetScore and available large-scale resources enabled identification of drugs that might potentially interact with metformin. Future validation experiments are necessary to explore the phenotypic outcome of interactions (i.e., both the positive and negative correlations) between metformin and other drugs.

## Discussion

Type II diabetes patients have been shown to suffer from increased risk of developing CRC (Berkovic *et al*., 2021). Mounting evidence suggests that the first line type II diabetes drug, metformin, could be beneficial in preventing the CRC development and improving the cancer prognosis (Berkovic *et al*., 2021). The MoA of metformin is pleiotropic, affecting both whole organism and cellular level (Pernicova and Korbonits, 2014; Rena *et al*., 2017), but not yet completely understood. Moreover, large-scale proteomic and phosphoproteomic studies have indicated that metformin does not affect a limited set of individual targets, but rather modulates the activity of multiple proteins and kinases resulting in an extensively rewired signaling network (Sacco *et al*., 2016). Due to the complexity of the metformin-rewired signaling network and the individual genomic variability, phosphoproteomics is a compelling approach to gain detailed mechanistic insights into the drug-induced signaling due to its ability to quantify thousands of P-sites across cell line panels. Additionally, recent technological, experimental, and computational advances increased the depth of such an analysis and facilitated the interpretation of the differential P-sites by inferring protein kinase activities and reconstructing signaling pathways mediated by these P-sites.

Here we performed a deep proteomic and phosphoproteomic analysis (**Figure 1**) of metformin-treated molecularly heterogenous CRC cell lines (Roumeliotis *et al*., 2017) after different timepoints to elucidate the MoA of metformin in CRC, address the heterogeneity of a drug response in a panel with a variable baseline proteome and phosphoproteome, and investigate the temporal dynamics of the cellular response to metformin. We confirmed the previous observation reported in a different cancer model that metformin remodeled the phosphoproteome within a 24-hour time window (Sacco *et al*., 2016). We also observed a phosphorylation upregulation for metformin-interacting proteins (Ma *et al*., 2022), which warrants future mechanistic investigation. Furthermore, we observed that the 24-hour response was highly heterogenous at multiple levels, as demonstrated specifically via 1) distinctive response at the level of individual differentially phosphorylated P-sites (**Figure 2**), 2) differential regulation of phosphorylation event occurring within specific biological processes (**Figure 2**), 3) differential kinase activity regulation (**Figure 5**), and 4) reconstructed signaling networks (**Figure 6**). These findings highlighted that the drug-induced signaling can be highly context-dependent in CRC and emphasized the importance of using multiple cell line models to study *in vitro* cancer cell signaling. The favorable reproducibility of Phos-DIA is fundamental to derive these observations in individual cell lines.

On the other hand, tens of phosphorylation sites and a set of protein kinases and biological processes were regulated in most cell lines (**Figure 4 & Table 1**). The identification of these phosphorylation events was facilitated by assigning each P-site a MetScore reflecting the robustness of a P-site regulation within the panel of CRC cell lines. MetScore enabled the stratification of P-sites into several segments for which we performed enrichment analysis to identify sequence motifs, associations with specific kinases, and biological functions. Moreover, we identified 55 most consistent P-sites regulated in at least 10 CRC cell lines which can be considered as the common “metformin signature” for CRC cells. The identification of a robust metformin signature might have the following clinical implications. First, in CRC patients with type II diabetes, metformin usage daily will have profound effects on the phosphoproteome of the tumor cells as demonstrated above. The metformin signature P-sites might serve as a resource to develop potential assays for monitoring metformin sensitivity and intended efficacy in cancer patients. Second, through regulation of specific P-sites, metformin might prime the cancer cells to be more sensitive or more resistant to conventional genotoxic chemotherapeutic agents or targeted cancer drugs. Thus, determination of how the phosphorylation signaling is rewired by metformin in cancer cells might be useful in optimizing cancer therapeutics.

Recently, there has been an increasing interest in repurposing of existing, well-characterized drugs for a novel usage including metformin (Fong and To, 2019). Metformin is an attractive repurposing drug due to its low cost and a compelling safety profile. Indeed, combinations of metformin with several genotoxic CRC chemotherapy drugs have been already reported (Khader *et al*., 2021; Liu *et al*., 2020; Richard and Martinez Marignac, 2015; Sang *et al*., 2020) indicating there is a potential for metformin in combination treatments. It would be ideal if metformin could either sensitize cancer cells to a drug leading to increased efficacy of the treatment, revert existing resistance, or decrease the deleterious side effects of chemotherapy. Since the synergies between drugs are rather rare and tend to be cell signaling dependent (Jaaks et al., 2022), we leveraged the identification of the metformin signature across the cell lines and queried established P-site abundance∼drug sensitivities correlations (Frejno *et al*., 2020). The rationale behind this approach relied on the assumption that if one or more P-site abundances are positively correlated with the drug sensitivity within the CRC65 panel (i.e., higher P-site abundance∼lower IC50), the upregulation of the P-site(s) might further sensitize the drug response and vice versa. Our preliminary results are promising regarding drug repurposing because most of the compounds identified have already been FDA-approved for other cancer indications (cetuximab, paclitaxel, pazopanib, nilotinib, and vorinostat) or in the late stages of clinical research development (navitoclax, SN-38, and YM-155); and for some of them (nilotinib, paclitaxel, pazopanib, and vorinostat), their synergy with metformin have been already reported in concrete literature in other cancer models. Our drug predictions are thus largely consistent with the existing experimental results. *In vitro* experiments are required to confirm metformin∼drug synergies in the context of CRC.

In conclusion, our study reveals novel and fundamental insights for understanding metformin-induced phosphorylation signaling, including the regulation timing and extent, the cell line dependency, and the signaling network and kinase landscape as the consequences of the regulation, providing a deep and substantial phosphoproteomic resource sharpening our views on metformin and its clinical potentials.

## Acknowledgements

YL thanks the support from the National Institute of General Medical Sciences (NIGMS), National Institutes of Health (NIH) through Grant R01GM137031 to YL. YL was also supported by a pilot grant from Cancer Systems Biology@Yale and a pilot grant from Yale Cancer Center at Yale University. We would like to thank Prem Subramaniam from the Department of Systems Biology at Columbia University for kindly providing the CRC cell lines.

## Author contributions

EG performed the cell culture experiment and sample processing. WL acquired the Orbitrap DIA data. BS, together with SMD (with the supervision of AD), performed the major data analysis. AD, GR, and JSR provided critical feedback to the manuscript. YL supervised and supported the study with help of JSR in data analysis. BS and YL wrote the paper.

## Material and Methods

### Cell culture and metformin treatment

The following 12 colorectal cancer (CRC) cell lines were used in this study: C2BBe1, COLO 205, HT115, LoVo, MDST8, NCI-H747, RKO, SNU-61, SW48, SW837, SW848, and T84. The cells were cultured on 10 cm dishes at 37 °C, humidified 5% CO^2^ in a complete medium containing L-glutamine, supplemented with 10% fetal bovine serum (Gibco, #26140079) and penicillin-streptomycin (Sigma-Aldrich, #P0781). The following cell culture media were used: Dulbecco’s Modified Eagle Medium (DMEM; Corning, #10-013-CV) was used for C2BBe1, HT115, and MDST8; RPMI-1640 (Thermo Scientific, #11875-093) was used for COLO 205, LoVo, NCI-H747, RKO, SNU-61, and SW48; Leibovitz’s L-15 Medium (Cytiva, #SH0525.01) was used for SW837 and SW948; DMEM/F12 (Thermo Scientific, #11330-032) was used for T84.

2 x 10e^6^ – 4 x 10e^6^ cells were seeded in a 10 cm dish, cultured for 24 h, and then treated with 10 mM metformin as described in a previous metformin phosphoproteomic study (Sacco *et al*., 2016). The following samples were harvested in three replicates: control samples at time points 0 and 24 h and metformin-treated samples at time points 30 min and 24 h. The 30-min time point was skipped for the total proteome analysis. After washing twice in PBS, the dishes were snap-frozen in liquid nitrogen for 2 min, and the cells were scraped into 200 µL of cell lysis buffer containing 10 M urea/ 100 mM ammonium bicarbonate, cOmplete™ protease inhibitor cocktail (Roche, #11697498001) and the Halt phosphatase inhibitors (Thermo Scientific, #78428). Cells were scraped using a cell scraper into a 2 mL Eppendorf tube. Tubes were vortexed for 30 s, snap-frozen in liquid nitrogen, and transferred to store at -80 °C until further sample processing. Triplicate dishes per cell line and condition were used as three whole process replicates for the analysis.

### Protein extraction and digestion

Cell pellets in lysis buffer were thawed and sonicated twice at 4 °C for 1 min using the VialTweeter device (Hielscher-Ultrasound Technology) (Liu et al., 2019) and centrifuged at 20,000 g for 1 hour to remove insoluble material. Protein concentration in the supernatant was determined using the Bio-Rad protein assay dye (Bio-Rad, #5000006). The protein samples (800 µg of protein per sample) were diluted in 6 M urea/100 mM ammonium bicarbonate buffer to 400 µl (final concentration = 2 µg/ µl), reduced by 10 mM DTT for 1 hour at 56 °C, and alkylated by 20 mM IAA in dark for 1 hour at room temperature (RT). The reduced and alkylated proteins were then subjected to a precipitation-based digestion as described previously (Collins et al., 2017). Briefly, five volumes of precooled precipitation solution containing 50% acetone, 50% ethanol, and 0.1% acetic acid were added to the protein mixture and kept at -20 °C overnight. The mixture was centrifuged at 20,000 x g for 40 min. The precipitated proteins were washed with precooled 100% acetone and centrifuged at 20,000 x g, 4 °C for 40 min. After acetone aspiration, the remaining acetone was evaporated in a SpeedVac. Next, 300 µL of 100 mM NH^4^HCO^3^ with sequencing grade porcine trypsin (Promega) at a ratio of 1: 20 were added and incubated overnight at 37 °C. The resulting peptide mixture was acidified with formic acid and then desalted with a C18 column (MarocoSpin Columns, NEST Group INC.) following the manufacturer’s instructions. The amount of the final peptides was determined by a nanodrop (Thermo Scientific).

### Phosphopeptide Enrichment

The phosphopeptide enrichment was performed using High-Select™ Fe-NTA kit (Thermo Scientific, #A32992) according to the kit instructions, as described previously (Gao et al., 2019). Briefly, the peptide-resin mixture was incubated for 30 min at room temperature while gently shaking and then transferred into a filter tip (TF-20-L-R-S, Axygen) to remove the supernatant (flow-through) by centrifugation. The resins were washed three times with 200 µL of washing buffer (80% I, 0.1% TFA) and twice with 200 µL of H^2^O. The phosphopeptides were eluted twice with 100 µL of elution buffer (50% ACN, 5% NH^3^•H^2^O) and dried in SpeedVac (Thermo Scientific). The amount of the final phosphopeptides was determined by a nanodrop (Thermo Scientific).

### Mass spectrometry measurements

For the LC-MS analysis, 1 µg of peptide and phosphopeptide mixture was used as described previously (Liu *et al*., 2019; Mehnert et al., 2019). The LC separation was using an EASY-nLC 1200 systems (Thermo Scientific) using a self-packed PicoFrit column (New Objective, Woburn, MA, USA; 75 µm × 50 cm length) self-packed with ReproSil-Pur 120A C18-Q 1.9 µm resin (Dr. Maisch GmbH, Ammerbuch, Germany). A 150-min measurement with buffer B (80% acetonitrile containing 0.1% formic acid) from 5% to 37% and corresponding buffer A (0.1% formic acid in H^2^O) during the gradient was used to elute peptides from the LC. The flow rate was set to300 nL/ min with the temperature controlled at 60 °C using a column oven (PRSO-V1, Sonation GmbH, Biberach, Germany). The Orbitrap Fusion Lumos Tribrid mass spectrometer (Thermo Scientific) was coupled with a NanoFlex ion source keeping the spray voltage at 2000 V and heating capillary at 275 °C. The DIA-MS method consisted of a MS1 survey scan and 33 MS2 scans of variable windows as described previously (Bruderer *et al*., 2017; Bruderer et al., 2019). The MS1 scan range is 350–1650 m/z and the MS1 resolution was 120,000 at m/z 200. The MS1 full scan AGC target value was set to be 2.0E6 and the maximum injection time was 100 ms. The MS2 resolution was set to 30,000 at m/z 200, and normalized HCD collision energy was 28%. The MS2 AGC was set to be 1.5E6 and the maximum injection time was 50 ms. The default peptide charge state was set to 2. Both of MS1 and MS2 spectra were recorded in a profile mode.

### Proteomic and phosphoproteomic DIA data analysis

DIA-MS data analyses for proteomics and phosphoproteomics were performed using Spectronaut v14 (Bruderer et al., 2015; Bruderer *et al*., 2017) using the library-free DirectDIA pipeline (Bruderer *et al*., 2017; Tsou *et al*., 2015) as described in detail in our previous study (Gao *et al*., 2021). The DIA runs were all directly searched against the Swiss-Prot protein database (September 2020, 20,375 entries). For the total proteome dataset, methionine oxidation and N-terminal acetylation were set as variable modifications, where carbamidomethylation at cysteine was set as a fixed modification. Additionally, for the searching of the phosphoproteomic dataset, phosphorylation at serine/threonine/tyrosine (S/T/Y) was enabled as a variable modification. Both peptide- and protein-FDR were controlled at 1%, and the resulting data matrix was filtered by “Qvalue”. The PTM localization function in Spectronaut v14 was enabled to localize and filter the phosphorylation sites using a PTM score > 0.75 (Bekker-Jensen *et al*., 2020; Rosenberger *et al*., 2017). All the other Spectronaut settings were kept as default, e.g., the ‘‘Interference Correction’’ was enabled, the ‘‘Global Normalization’’ (‘‘Median’’) was used, the quantification was performed at the MS2 level using peak areas, and the Top 3 peptide precursors (‘‘Min: 1 and Max: 3’’) were averaged (mean) for representing protein quantities in all DIA analyses.

### Data processing

For the total proteome analysis, the protein pivot reported was exported from Spectronaut. Relative intensity values < 500 were replaced by NA, the data were log2 transformed and normalized using LOESS (Smyth, 2005). For the phosphoproteome data analysis, phosphopeptide precursor level pivot report was exported from Spectronaut using two different localization probability score cutoffs (0.75 and 0) using a strategy described in our previous work (Gao *et al*., 2021). The first report (localization probability > 0.75, class I sites (Olsen *et al*., 2006), n = 117,067 unique modified peptide precursor ions) was used to identify phosphopeptide precursor ids with confidently localized P-sites in at least one sample. The second report (localization probability > 0, n = 132,887 unique modified peptide precursor ions) was filtered to only contain the 117,067 sites from the first report and was used to extract the phosphopeptide precursor intensity values. The resulting total number of identified unique phosphoprecursors was 92,422. To select the most representative phosphopeptide precursor per unique phosphopeptide id, “**phos.id**”, we applied the following filtering criteria: i) the precursor with the most valid values across the 144 MS runs, ii) in case there were multiple precursor ids with the same number of valid values, the top intensity id was used (sum intensity across samples). The number of resulting unique phos.ids was 64,680 from 7,250 phosphoproteins, these ids preserved information about multiply phosphorylated peptides and were used for the downstream analysis. These ids corresponded to 56,080 unique P-sites (i.e., protein + site localization, “**phos.id.exp**”), these were used to perform the data annotation for all site-specific analyses. Phosphopeptide intensities were filtered to remove all intensities < 500, log2 transformed, and normalized using LOESS (Smyth, 2005).

### Detection of net phosphorylation changes

When specified, the relative protein abundances were regressed out from the respective relative phosphopeptide abundance values to detect net phosphorylation changes using linear regression as described previously (Roumeliotis *et al*., 2017). Briefly, phosphopeptide intensities were set as the dependent variables (y) and matched protein intensities were set as the independent variables (x). The residuals of the y∼x linear model were used as the phosphorylation levels not driven by protein abundance levels.

### Consensus clustering

Consensus clustering (Monti et al., 2003; Wilkerson and Hayes, 2010) was performed with the ConsensusClusterPlus R package (Wilkerson and Hayes, 2010) using log2 fold changes (metformin 24 hours / control 24 hours) or normalized log2 intensities (“steady-state”). Only those ids with quantitative values in all cell lines were used and the top 30% most variable ids were selected based on their variability across samples (median absolute deviation; MAD). The number of clusters selected for metformin response classification was four.

### Statistical analysis

Statistical analysis to identify differentially abundant proteins and phosphopeptides was performed in Perseus v1.6.14.0 (Tyanova et al., 2016) using a two-sided Student’s t-test for every comparison with at least 2 valid values per each compared group to avoid missing values imputation. Protein groups and phos.ids with p < 0.01 and a fold-change > 1.5 were reported as significant. To identify differentially abundant phos.ids at 30 minutes and 24 hours, the control samples at 0 hours and 24 hours were used, respectively, to consider the effect of nutrient exhaustion in media on the phosphoproteome after 24 hours of cultivation. To identify differentially abundant proteins after 24 hours, we used the matched 24-hour control samples as well.

### MetScore calculation

To identify phos.ids consistently responding to metformin treatment across the 12 cell lines, we developed a scoring system which we termed MetScore. After statistical analysis (p < 0.01 and fold-change > 1.5; two-sided Student’s *t*-test) only those ids subjected to statistical analysis in all cell lines (i.e., having required number of valid values as described above) were used leading to a complete matrix containing 14,032 unique phos.ids for the 24-hour dataset. The score was calculated across the 12 cell lines, i.e., when a phos.id was significantly upregulated, we added one (+1) to the score, and when an id was significantly downregulated, we subtracted one (-1) from the resulting score leading to a score distribution from -12 to 12 reflecting the consistency of the response across the cell lines. Based on the MetScore, the phos.ids were stratified into five segments for the downstream enrichment analysis: MetScore G1 (MetScore ɛ [4, 12], n = 1,207, 8.6% of the dataset), MetScore G2 (MetScore ɛ [3, 2], n = 1,660, 11.83%), MetScore G3 (MetScore ɛ [-1, 1], n = 8,344, 59.86%), MetScore G4 (MetScore ɛ [-2, - 3], n = 1,607, 11.45%), and MetScore G5 (MetScore ɛ [-4. -12], n = 1,214, 8.65%).

### Sequence analyses

To compare the MetScore G1 and G5 P-sites sequence windows (**Figure 4D**), we used iceLogo (Colaert *et al*., 2009) with p < 0.05 cutoff. To extract enriched sequence motifs (**Figure 4E**), we employed motifeR (Wang *et al*., 2019) with a minimal occurrence of a motif set to 20 and the motif enrichment p-value < 0.000001.

### Data annotation and functional enrichment analyses

Protein-specific annotation was downloaded from DAVID Bioinformatics Resources 6.8 [https://david.ncifcrf.gov/] (Huang da et al., 2009; Huang da et al., 2008; Sherman et al., 2022). Site-specific annotation was downloaded from the PhosphoSitePlus database [www.phosphosite.org] (Hornbeck *et al*., 2012; Hornbeck *et al*., 2015), PTMsigDB v1.9.0 (Krug *et al*., 2019), and the OmniPath database (Türei *et al*., 2016; Türei *et al*., 2021). The validated lysosomal metformin-interacting protein list used in **Figure 3** (n = 113) was downloaded from the recent Ma et al. publication (Ma *et al*., 2022). The functional score used in **Figure 4** was downloaded from the Ochoa et al. publication (Ochoa *et al*., 2020). The 1D enrichment and Fisher’s exact test analyses were performed in Perseus v1.6.14.0 (Tyanova *et al*., 2016) or in R (R Core Team, 2020). For the protein-specific annotation enrichment analyses of the phosphoproteome dataset, the analyses were performed relative and not relative to unique protein ids to include the information about the number of phosphorylation events per protein in a pathway but also consider the number of unique protein ids identified per pathway. For the 1D enrichment analyses using fold changes between conditions (such as in **Figure 2E**), a looser filtering was applied to the data. First, the replicate values were averaged (mean) for each sample requiring at least one value per triplicate, and then a fold-change was calculated between conditions leading to a more complete matrix containing more sites for a more sensitive biological analysis.

### Kinase activity estimation and signaling network modeling

After data processing and normalization, differential analysis was performed between each control and metformin-treated cell line after 24 hours using the limma R-package (Ritchie et al., 2015). The standard sequence of lmFit, contrasts.fit, and eBayes was used, and the limma results were corrected for multiple testing using FDR correction (Benjamini-Hochberg method). For each cell line, kinase activity was estimated based on the abundance of the direct target P-sites. Post translational modifications were extracted from OmniPath (Türei *et al*., 2016; Türei *et al*., 2021) and filtered to keep only phosphorylation and dephosphorylation events. Interactions reported exclusively in the ProtMapper database were removed after noticing inconsistent interactions leading to a total of 29,445 signed kinase-P-site interactions of 580 different kinases. Additionally, t-values from the differential analysis of each P-site after metformin stimulation using the limma R-package were passed to the run_viper function from the decoupleR R-package (Alvarez et al., 2016; Badia-i-Mompel *et al*., 2022). Only kinases with at least five measured targets were included in the kinase activity estimation.

Signaling networks were contextualized for each cell line using the R-package PHONEMeS (Terfve *et al*., 2015) (PHOsphorylation NEtworks for Mass Spectrometry) by combining the information from large-scale MS phosphoproteomic data with prior knowledge of signaling. Based on integer linear programming (ILP) implementation for causal reasoning, PHONEMeS finds a path that connects deregulated P-sites with deregulated kinases in a coherent manner. For that a selection of deregulated P-sites and deregulated kinases and a prior knowledge network (PKN) are required. The prior knowledge network contains protein-protein and kinase-P-site interactions retrieved from OmniPath, resulting in a network with a total of 49,219 edges and 18,966 unique nodes. The top 15% of P-sites based on their absolute t-value were selected as deregulated and their t-values were passed to PHONEMeS. For the selection of deregulated kinases, the top 15% based on their absolute normalized enrichment score (NES) were chosen and selected as up-regulated (1) with a NES over zero and down-regulated (-1) with a NES lower than zero. For cell lines where AMPK (PRKAA1) belonged to the top 15% based on its absolute NES, this node was manually removed from the selected kinases, to be able to gain insights about its upstream regulation. Additionally, kinases with an absolute NES lower than 0.5 were passed to PHONEMeS to be removed from the PKN. Before connecting deregulated P-sites to kinases, the PKN was pruned by removing all nodes that are not connected 50 steps downstream and upstream of the selected kinases and P-sites, respectively. For each node in the resulting networks the inferred activity of PHONEMeS was compared to the estimated activity from decoupleR if possible. If these activities did not agree on their direction of deregulation the nodes were either removed from the PKN (if NES < 2) or added to the inputted deregulated kinases (if NES ≥ 2). PHONEMeS was re-run with the adapted inputs and all steps were repeated as described above until a network solution was found where the PHONEMES inferred and decoupleR estimated activities were coherent for all overlapping nodes. Protein-protein networks were constructed for each cell line by removing interactions between kinase and phosphorylation sites from the network. To focus on regulation surrounding PRKAA1, subnetworks were extracted containing all nodes and edges two steps up- and downstream of PRKAA1. For the comparison of subnetworks between different cell lines, a general backbone was created containing the nodes of all subnetworks, with cell line-specific nodes highlighted.

### Drug signature comparison

To identify drugs potentially interacting with metformin CRC, we queried the ATLANTiC database (Frejno *et al*., 2020) [http://atlantic.proteomics.wzw.tum.de] for the 55 metformin signature sites (absolute MetScore ≥ 10). For each site, we extracted the top 50 most significant drug associations based on “Simple correlation” analysis *p*-value. Using these drug∼P-sites associations, we generated a network and finally selected drugs shown in **Figure 7** based on their degree (number of edges). The correlation analysis presented in the Supplementary material was performed as follows, similarly to a previous study (Zhang et al., 2021). Drug signature comparisons were performed by querying Touchstone-P, a library of phosphoproteomic signatures of the relative abundances of approximately 100 phosphorylation sites (P100) (Abelin *et al*., 2016) from a panel of 7 different cell lines treated with 118 small-molecule drugs (Dele-Oni *et al*., 2021; Litichevskiy *et al*., 2018) available through the ConnectivityMap web interface (http://clue.io/proteomics-query). The input P100 phosphorylation signatures of the 12 CRC cell lines treated with metformin for 24 hours were compared to each signature in the library a connectivity score ranging from -1 (strong negative connection/most “opposite” profile) to 1 (strong positive connection/most similar profile) was reported for further analysis. The second analysis was performed using PhosFate Profiler (Ochoa *et al*., 2016) [http://phosfate.com]. Briefly, the relative log2 fold changes induced by metformin in individual cell lines were submitted to retrieve the correlation of the signaling response with 399 previously published phosphoproteomes.

### Data visualization

Most data visualization was performed in R (R Core Team, 2020). The following R packages were used to visualize the data: “ggplot2” (boxplots, violin plots, volcano plots, and histograms), “plot” (principal component analysis), “pheatmap” (heatmaps), “corrplot” (correlation plots), and “UpsetR” (UpSet plots). Barplots were generated using using GraphPad Prism version 8.0 (GraphPad Software, San Diego, California USA). The kinome trees were generated using Coral (Metz et al., 2018). All networks were visualized in Cytoscape v3.9.1 (Shannon et al., 2003).

### Data availability

Raw data will be available upon manuscript publication.

## Supplementary figure legends

**Supplementary Figure 1: Correlation analysis of the total proteome data (related to Figure 1).** Pearson correlation between samples was calculated using log2-transformed protein abundances. The heatmap shows hierarchical clustering of the correlation coefficients between samples.

**Supplementary Figure 2: Correlation analysis of the phosphoproteome data (related to Figure 1).** Pearson correlation between samples was calculated using log2-transformed phosphopeptide abundances. The heatmap shows hierarchical clustering of the correlation coefficients between samples.

**Supplementary Figure 3: Phosphopeptide level change was not driven by protein abundance change after 24 hours of metformin treatment (related to Figure 2). (A)** The relative protein abundances were regressed out from the respective relative phosphopeptide abundance values to detect net phosphorylation changes using linear regression (see **Methods**). The scatter plot shows the phosphopeptide-specific Pearson correlation of the phosphopeptide abundance and abundance of the corresponding total protein (phos∼prot correlation) before (x axis) and after (y axis) the regression. **(B)** Pearson correlation coefficients of the log2 metformin/control fold change prior and after the regression. Each dot represents a value of an individual cell line. The red dot represents the median value. **(C)** Examples of the log2 metformin/control fold change prior and after the regression in four cell lines.

**Supplementary Figure 4: Cell line-specificity of the phosphoproteome level response at the individual sites level (related to Figure 2). (A-B)** UpSet plots show the overlaps of the significantly up- **(A)** or down- **(B)** regulated phosphopeptides between individual cell lines.

**Supplementary Figure 5: Consensus clustering analysis of the metformin response (related to Figure 2). (A)** Four clusters revealed by the ConsusClusterPlus algorithm based on the top 30% of most variable metformin response (log2 fold change metformin/control after 24 hours) across the 12 cell lines. **(B)** Clustering based on metformin response mostly does not resemble clusters retrieved based on the steady-state (control) proteome and phosphoproteome.

**Supplementary Figure 6: PHONEMeS reconstructed signaling networks in individual cell lines (related to Figure 6). (A-L)** The border color corresponds to the protein activity scores calculated using PHONEMeS; the fill color mapping illustrated the decoupleR kinase activity score. The shape indicates whether the node is a P-site measured in our data (ellipse), kinase differentially perturbed in our data (diamond), or protein inferred by the algorithm to be a part of the signaling network (rectangle). The size corresponds to the number of out-going edges.

**Supplementary Figure 7: Metformin∼drug profile correlation analysis (related to Figure 7). (A)** Heatmap of all metformin∼drug profile correlations (normalized correlation score) based on the P100 reduced representative phosphosignature dataset. **(B)** Heatmap of all correlations (Pearson correlation) of metformin with paclitaxel treatment datasets based on the PhosFate profiler webtool. **(C)** Drugs associated the most with the non-G3 MetScore P-sites based on the PTMsigDB.

## Supplementary table legends

**Supplementary Table 1: Overview of the phosphoproteome relative quantification results.** Statistical analysis results and a MetScore after 24 hours of metformin treatment are provided.

## Abbreviations

AMPK: 5’ AMP-activated protein kinase
CAMKK2: calcium/calmodulin-dependent protein kinase kinase 2/beta
CDKs: cyclin-dependent kinases
CRC: colorectal cancer
DIA-MS: data-independent acquisition mass spectrometry
FDR: False discovery rate
LC: liquid chromatography
LKB1/STK11: serine/threonine-protein kinase STK11
MAPK3K7/TAK1: mitogen-activated protein kinase kinase kinase 7
MAPKs: mitogen-activated protein kinases
MoA: mechanisms of action
NES: normalized enrichment score
PER2: Period circadian protein homolog 2
PGRMC2: Progesterone receptor membrane component 2
PKN: prior knowledge network
mTOR: serine/threonine-protein kinase mTOR
MS: mass spectrometry
P-site: phosphorylation site
T2D: type 2 diabetes

