## Supplementary Figures 1-7 for "Deep Phosphoproteomic Elucidation of Metformin-Signaling in Heterogenous Colorectal Cancer Cells"

### Supplementary figure 1

#### Total proteome abundance correlation matrix

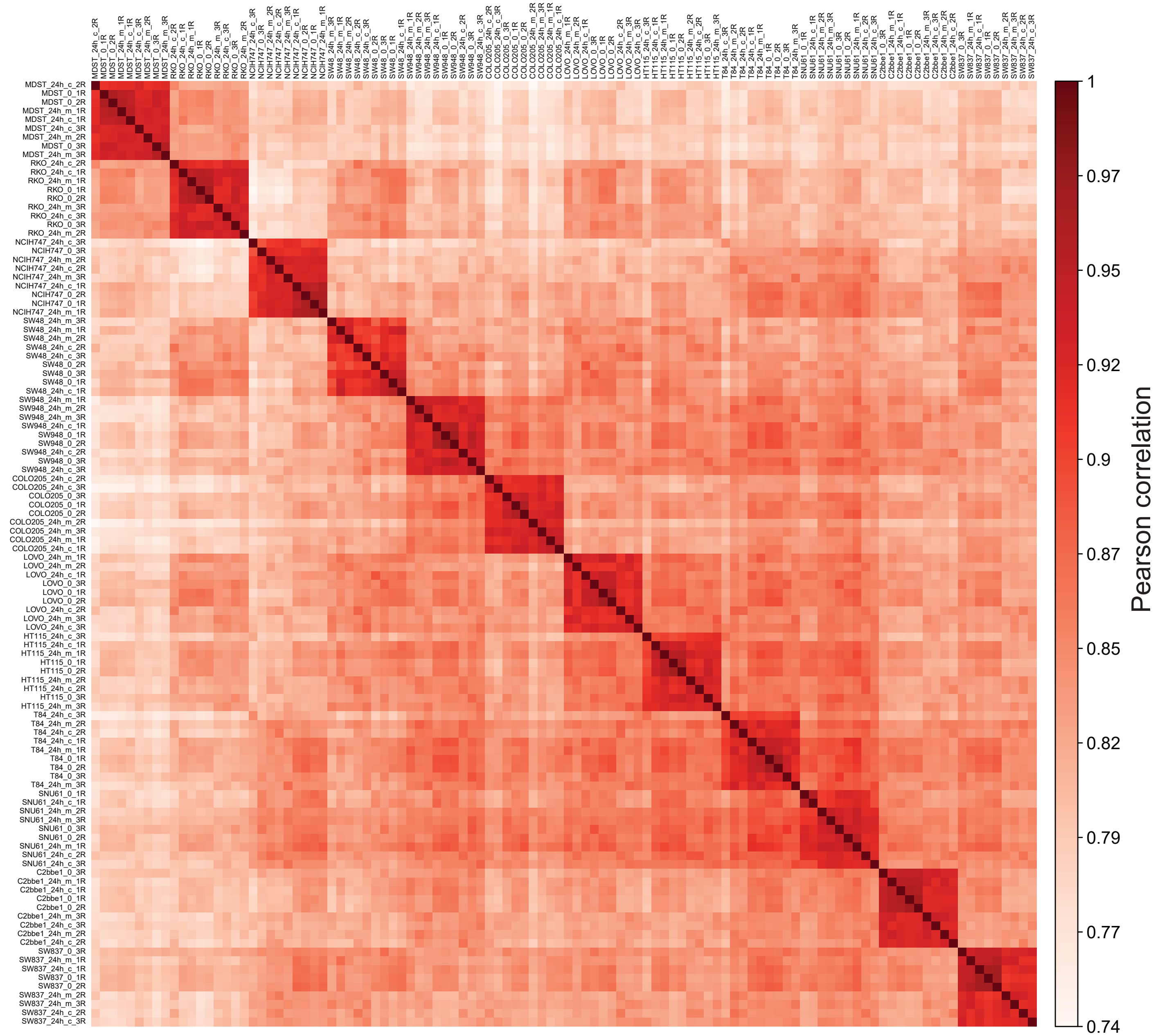

#### Supplementary figure 2

##### Phosphoproteome abundance correlation matrix

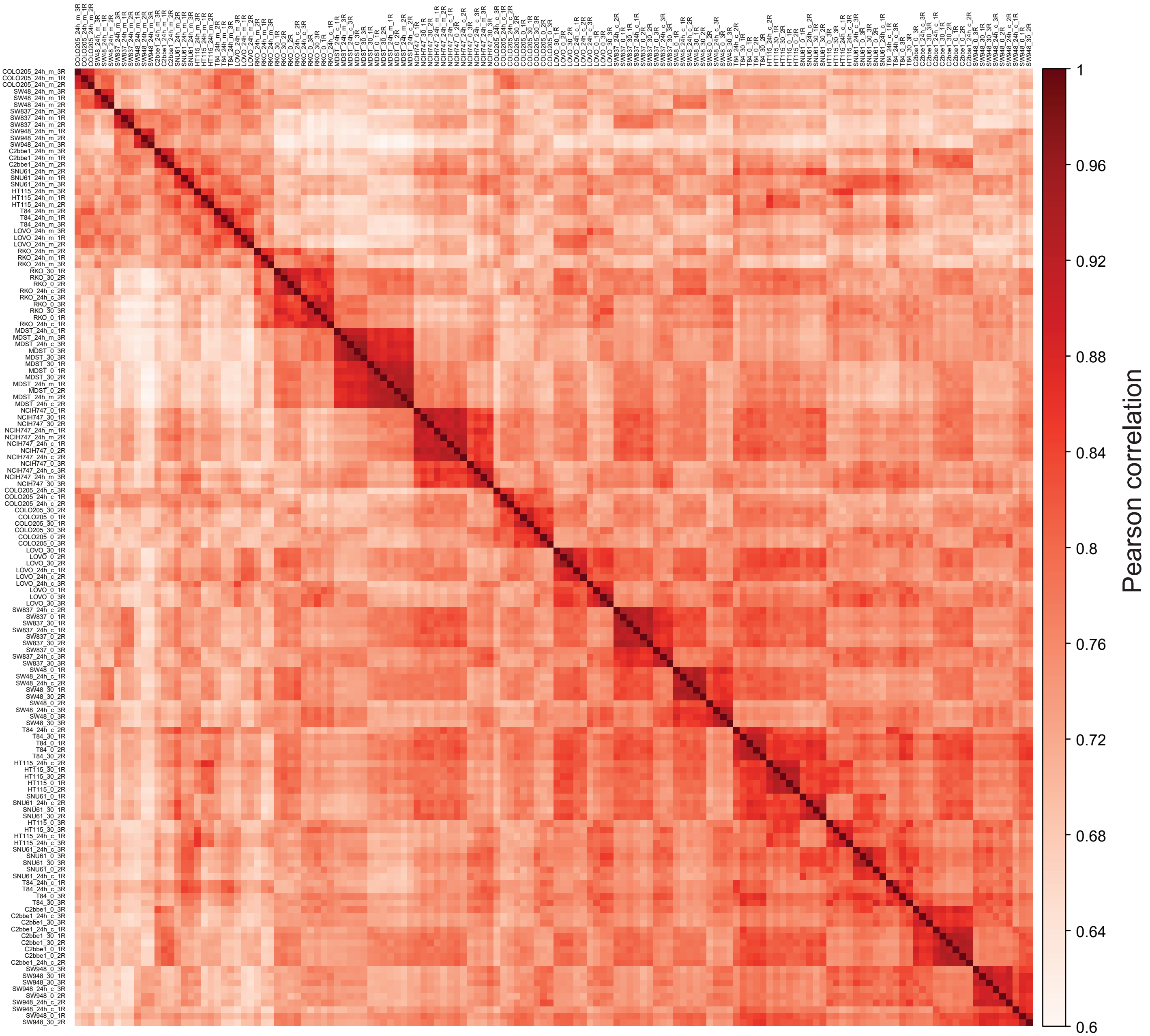

#### Supplementary Figure 3

A

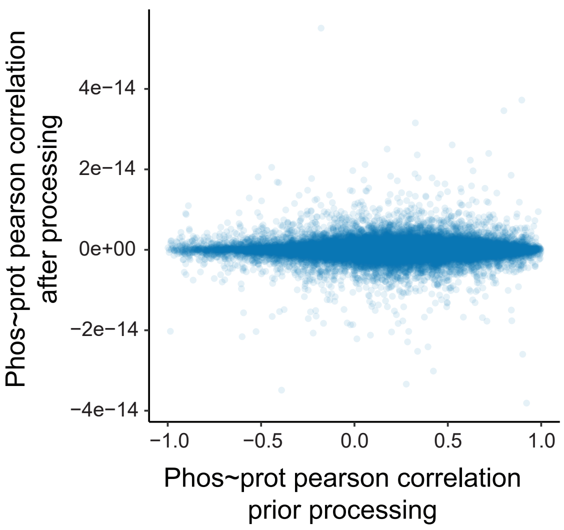


B

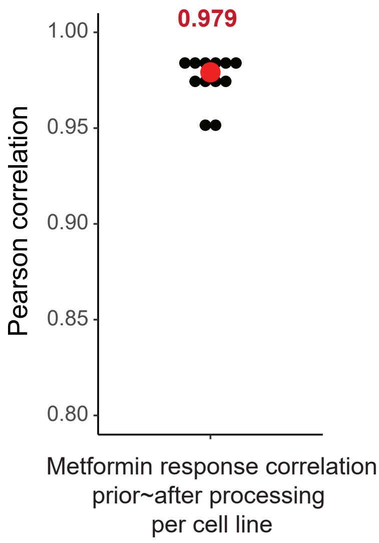


C

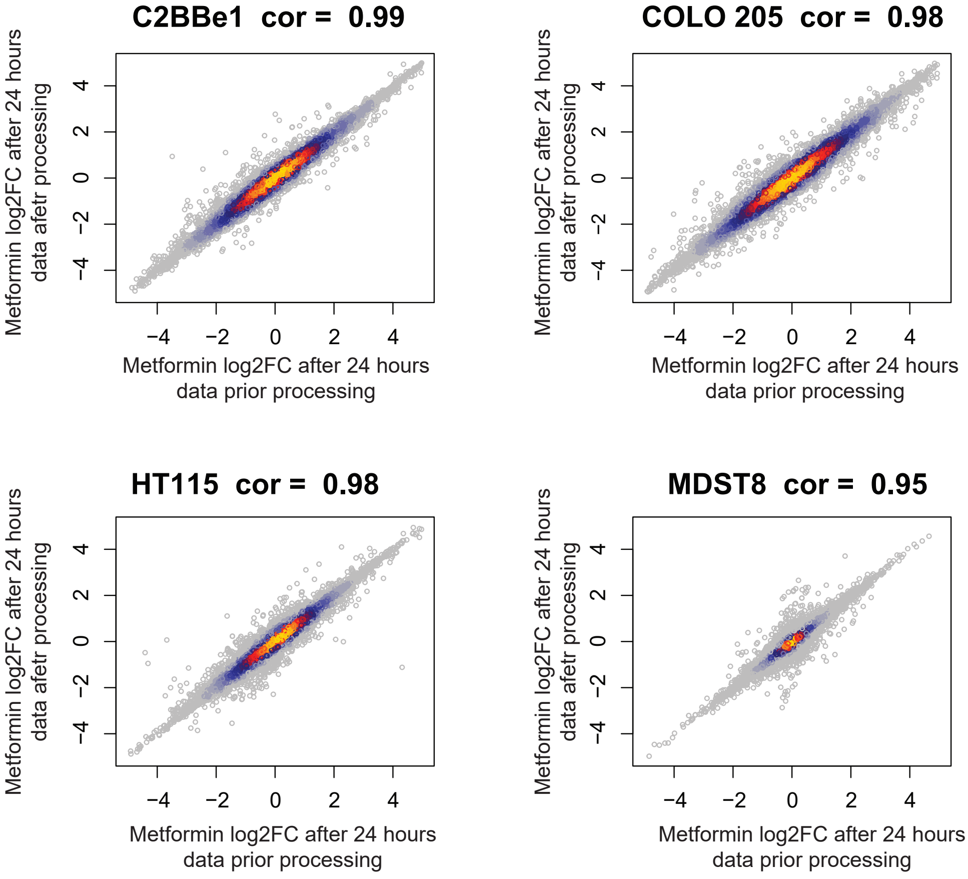

### Supplementary Figure 4

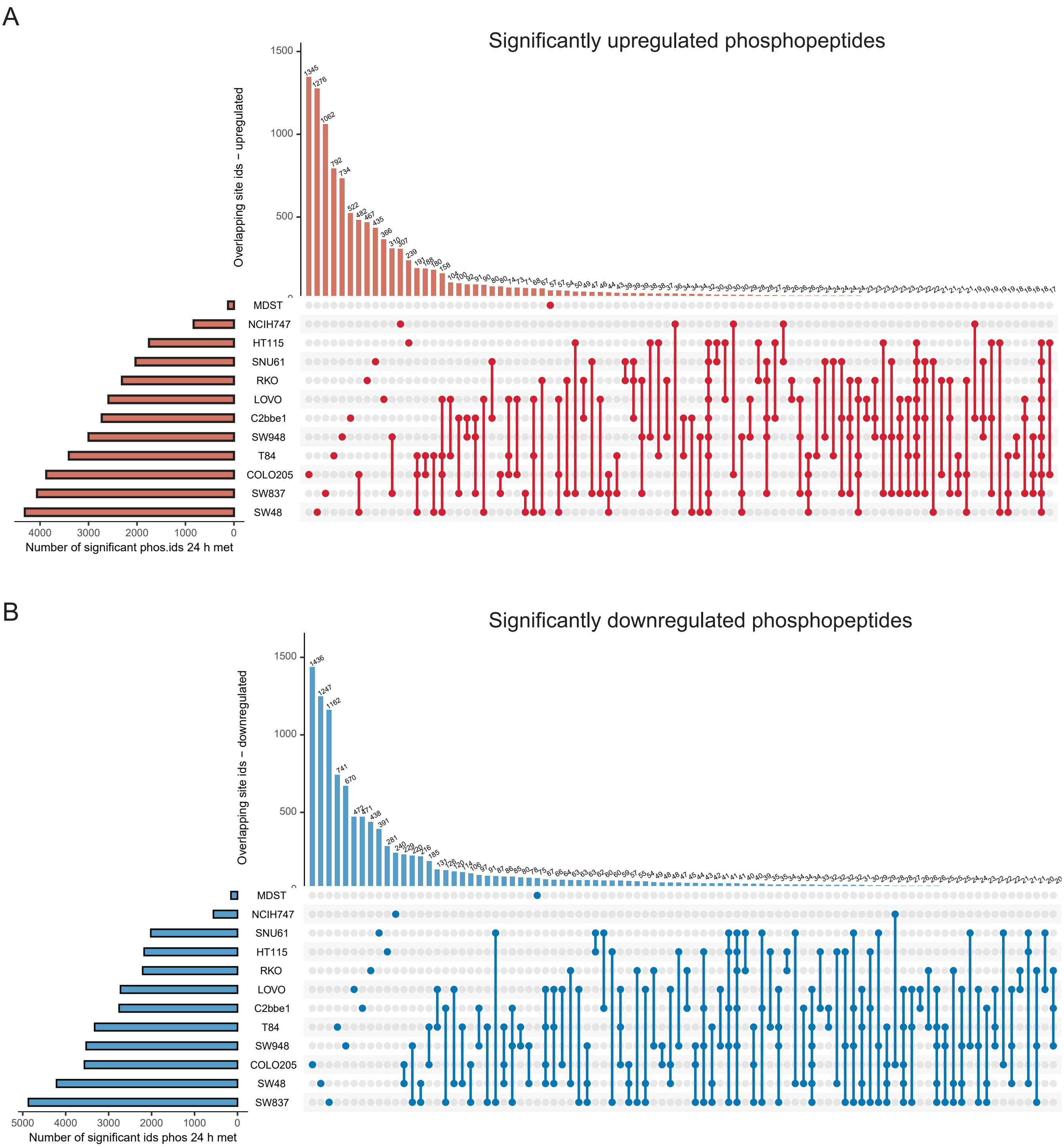

### Supplementary Figure 5

A

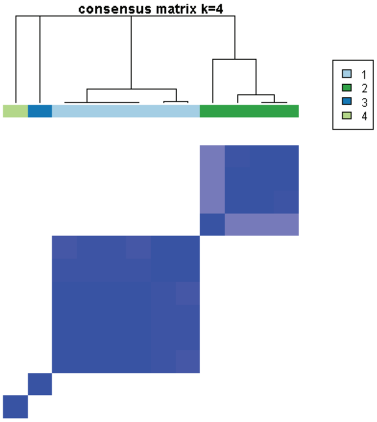


B

| Cell Line | Metformin response 24 hours phosphoproteome | Steady-state total phosphoproteome | Steady-state total proteome | Proteomic cluster Roumeliotis, 2017 |
|---|---|---|---|---|
| C2bbe | 1 | 1 | 1 | Cluster 2 |
| COLO205 | 2 | 2 | 2 | Cluster 3 |
| HT115 | 1 | 3 | 3 | Cluster 2 |
| LOVO | 2 | 4 | 3 | Cluster 1 |
| MDST8 | 3 | 1 | 3 | Cluster 4 |
| NCIH747 | 4 | 1 | 1 | Cluster 5 |
| RKO | 1 | 4 | 3 | Cluster 1 |
| SNU61 | 1 | 3 | 1 | Cluster 2 |
| SW48 | 2 | 4 | 3 | Cluster 1 |
| SW837 | 1 | 1 | 4 | Cluster 5 |
| SW948 | 1 | 2 | 2 | Cluster 2 |
| T84 | 2 | 1 | 1 | Cluster 2 |

Supplementary Figure 6A

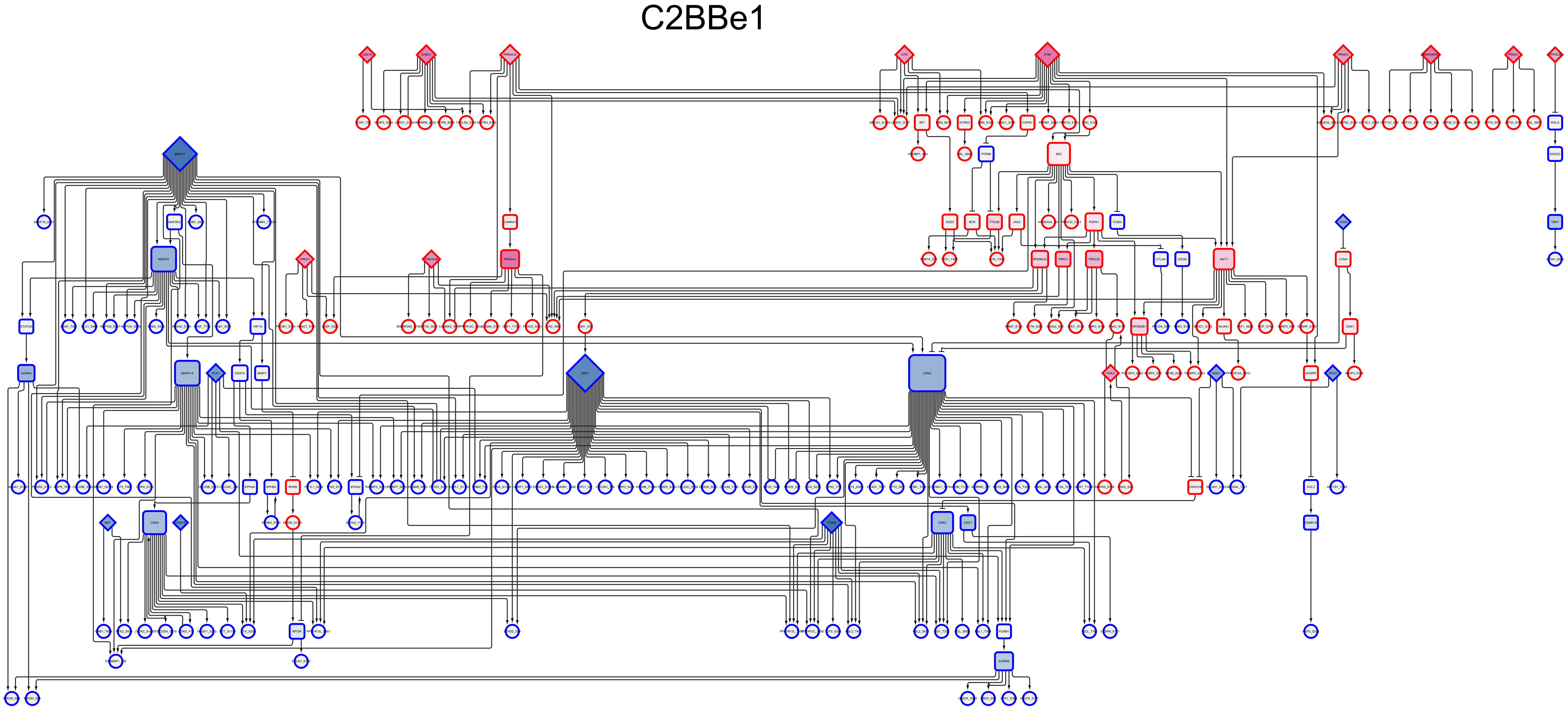

Supplementary Figure 6B

### COLO 205

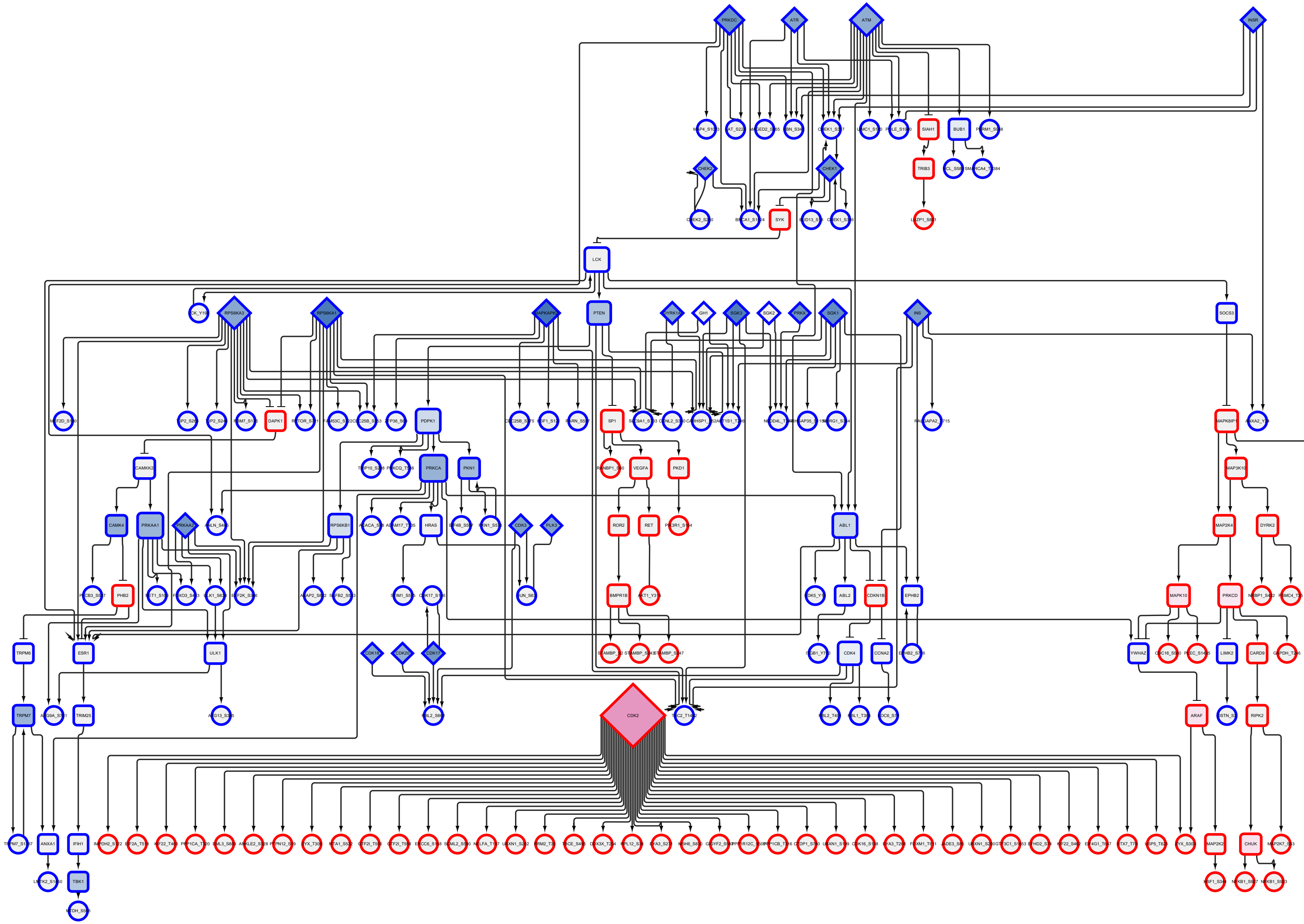

Supplementary Figure 6C

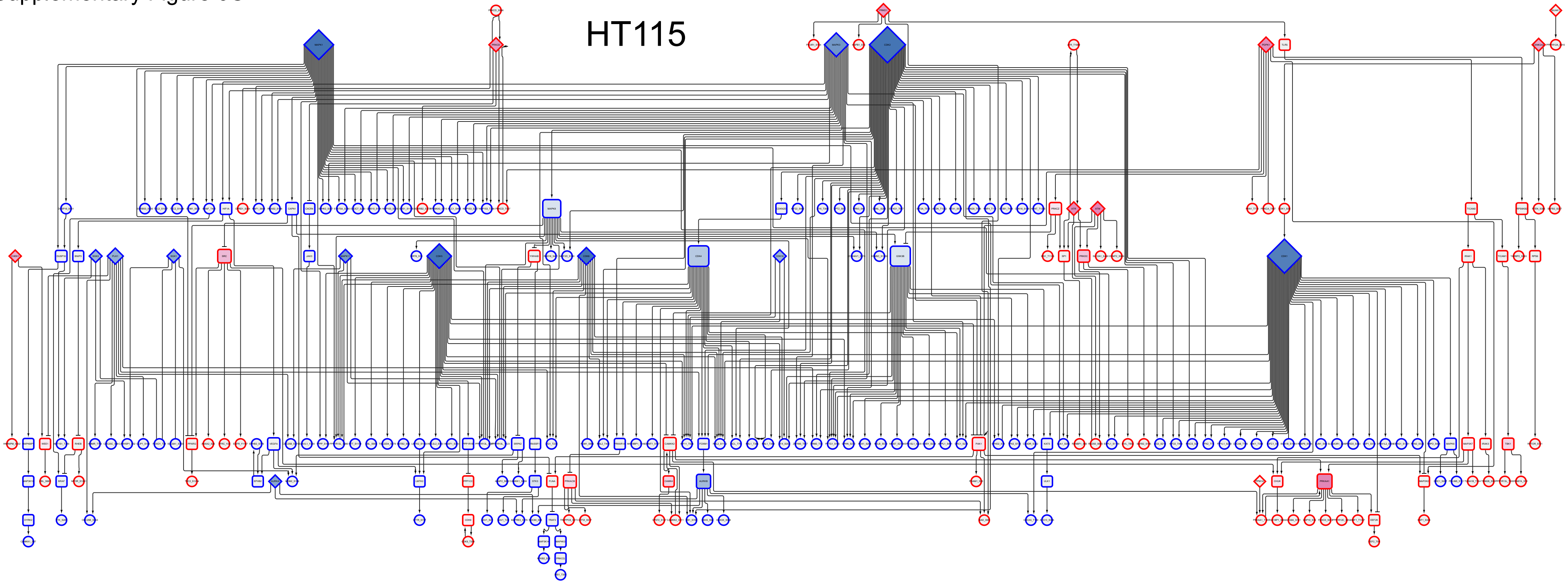

Supplementary Figure 6D

### LoVo

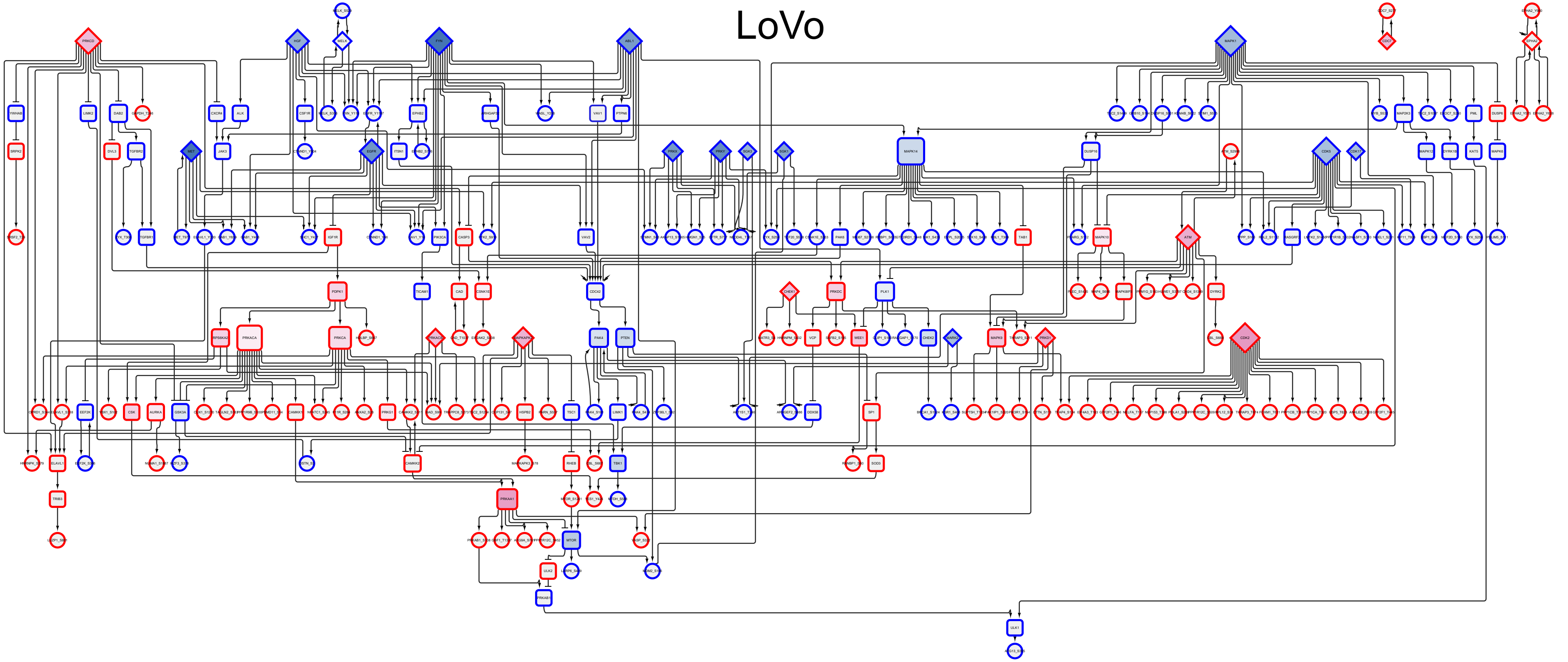

Supplementary Figure 6E

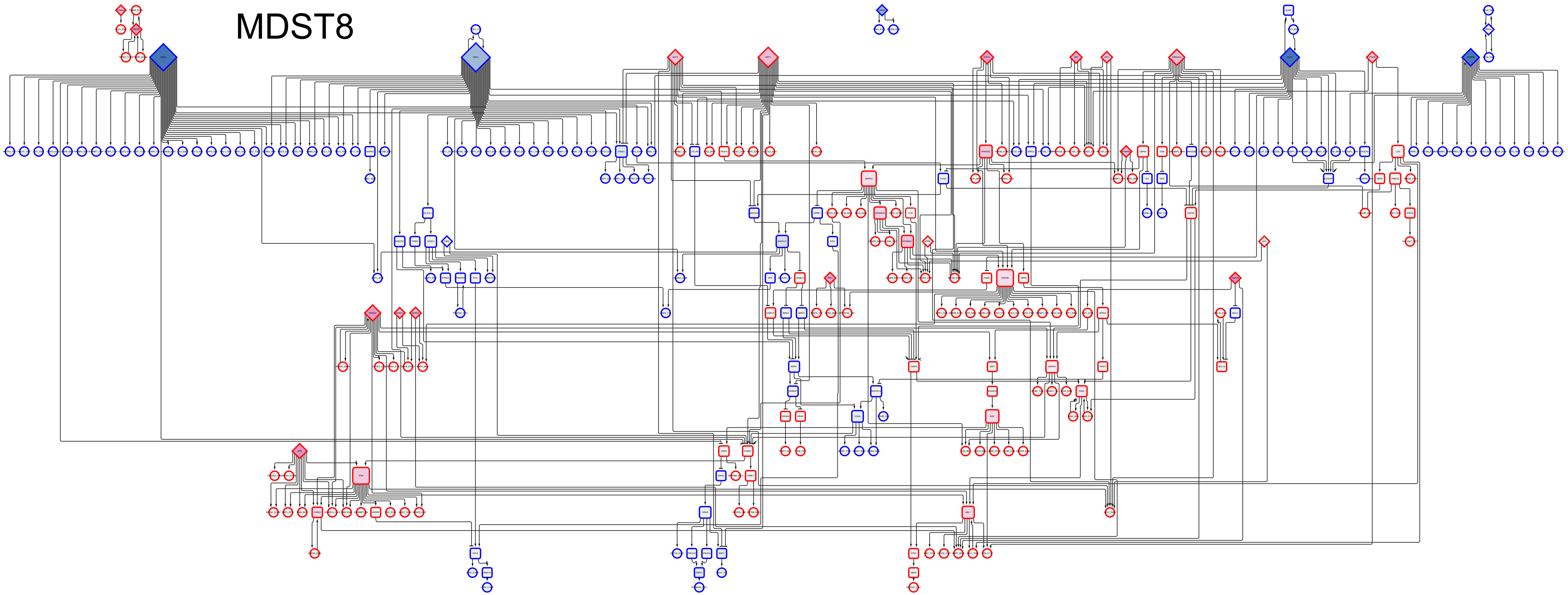

Supplementary Figure 6F

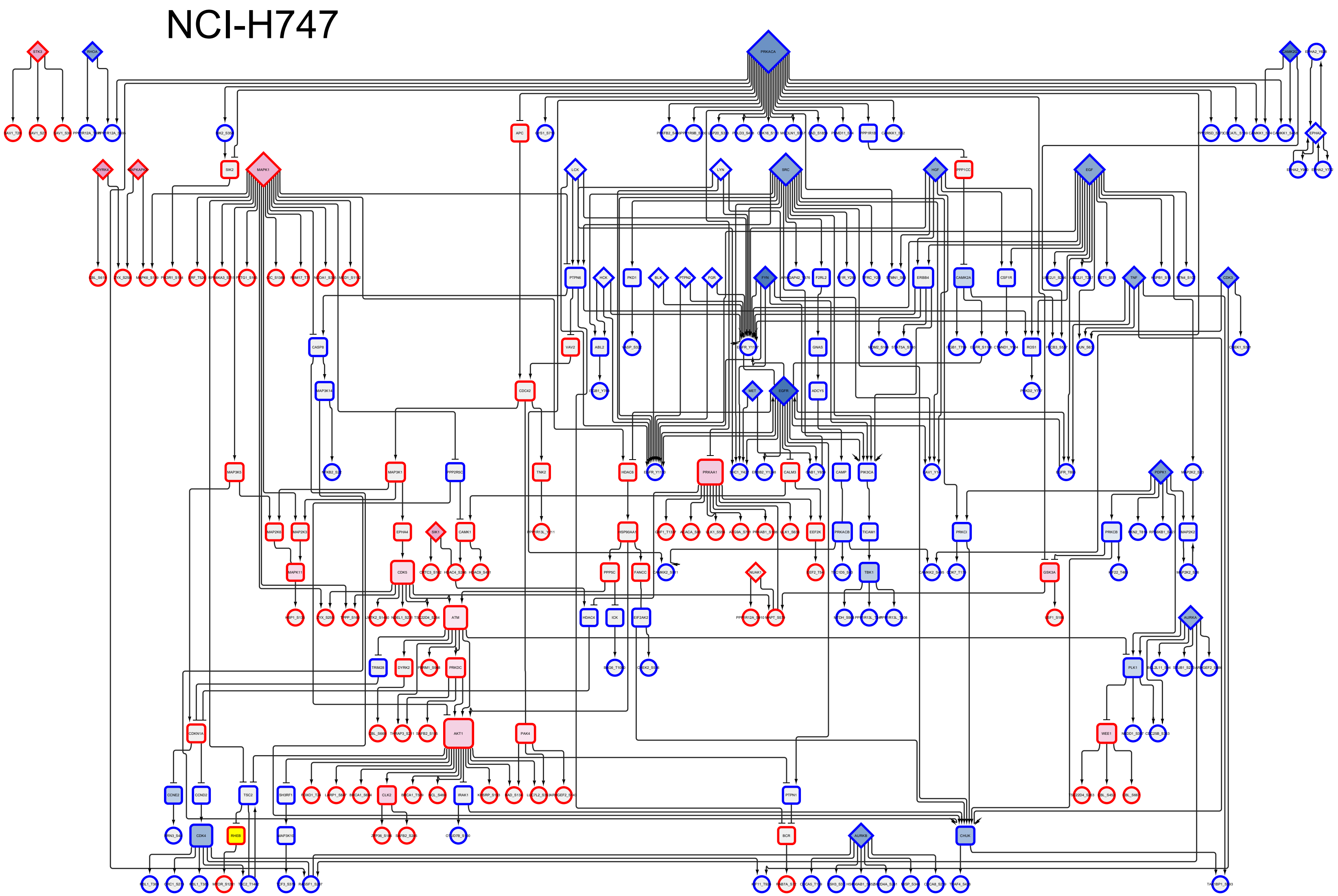

Supplementary Figure 6G

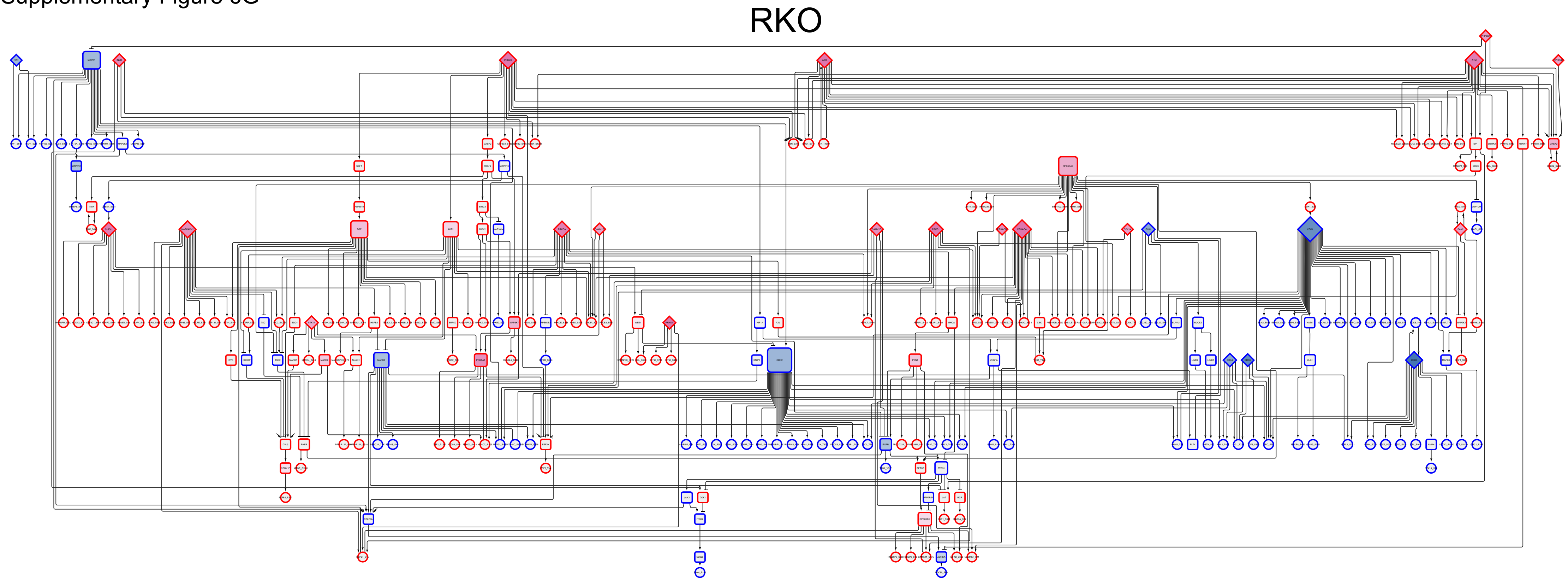

Supplementary Figure 6H

### SNU-61

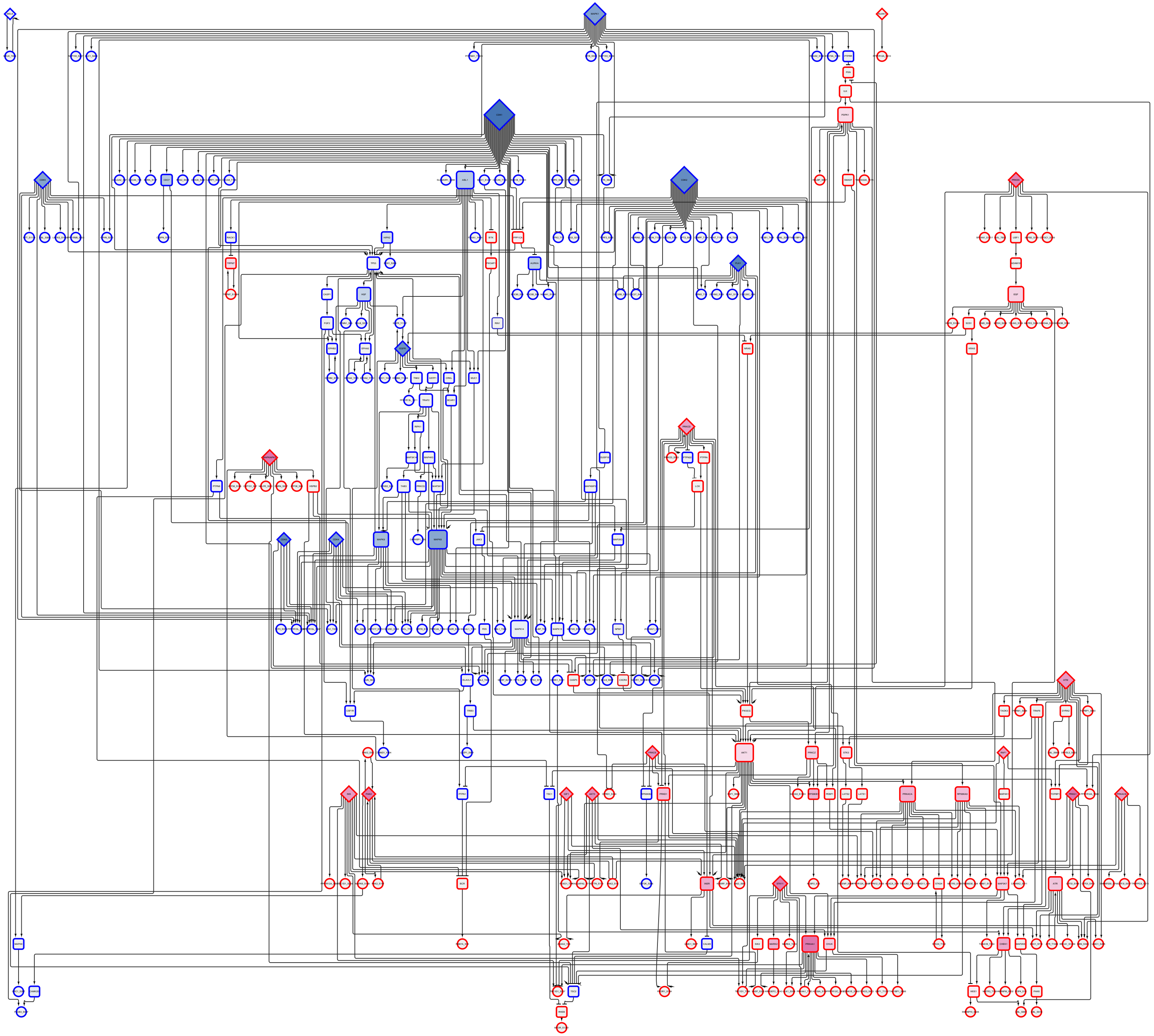

Supplementary Figure 6I

# SW48

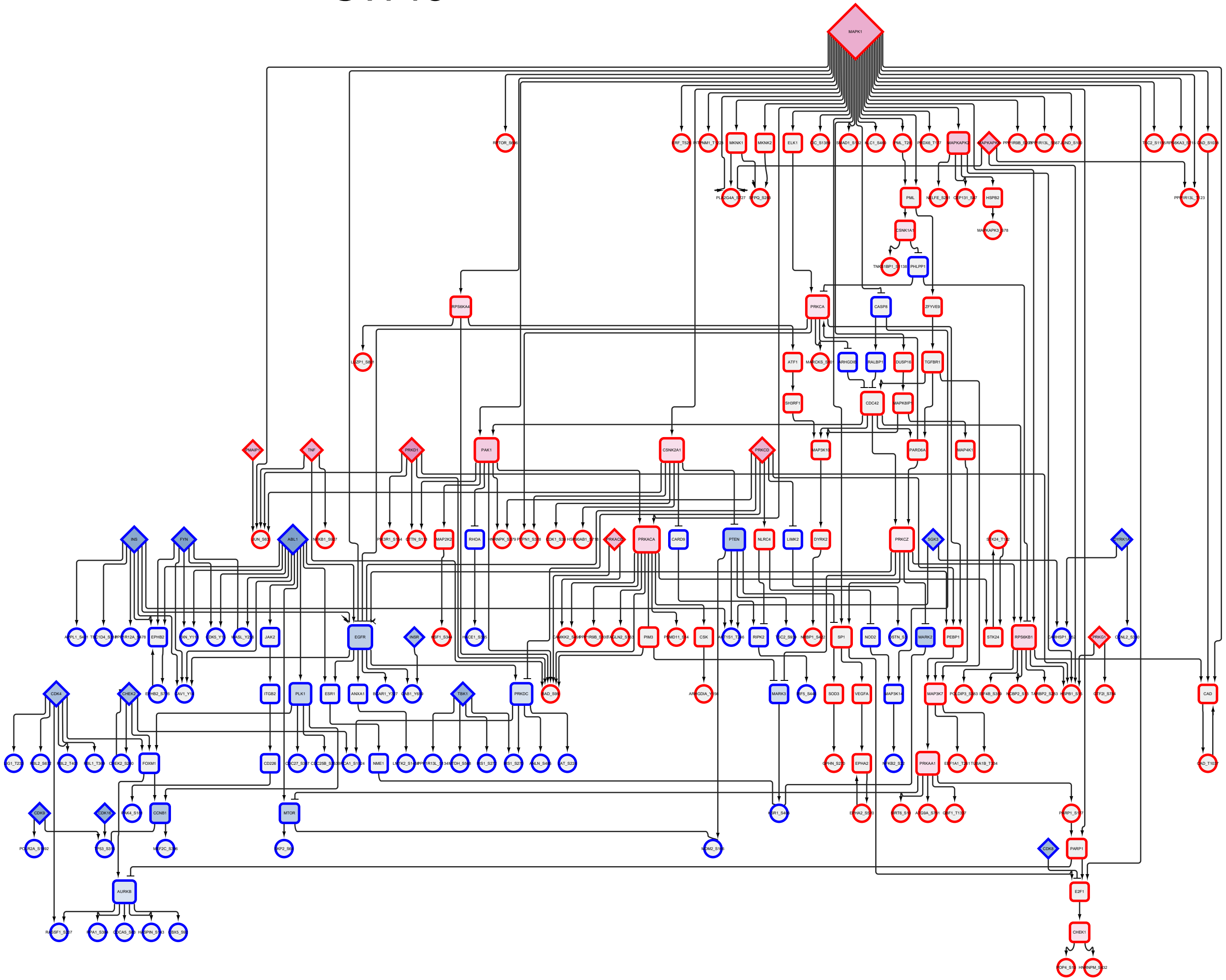

Supplementary Figure 6J

# SW837

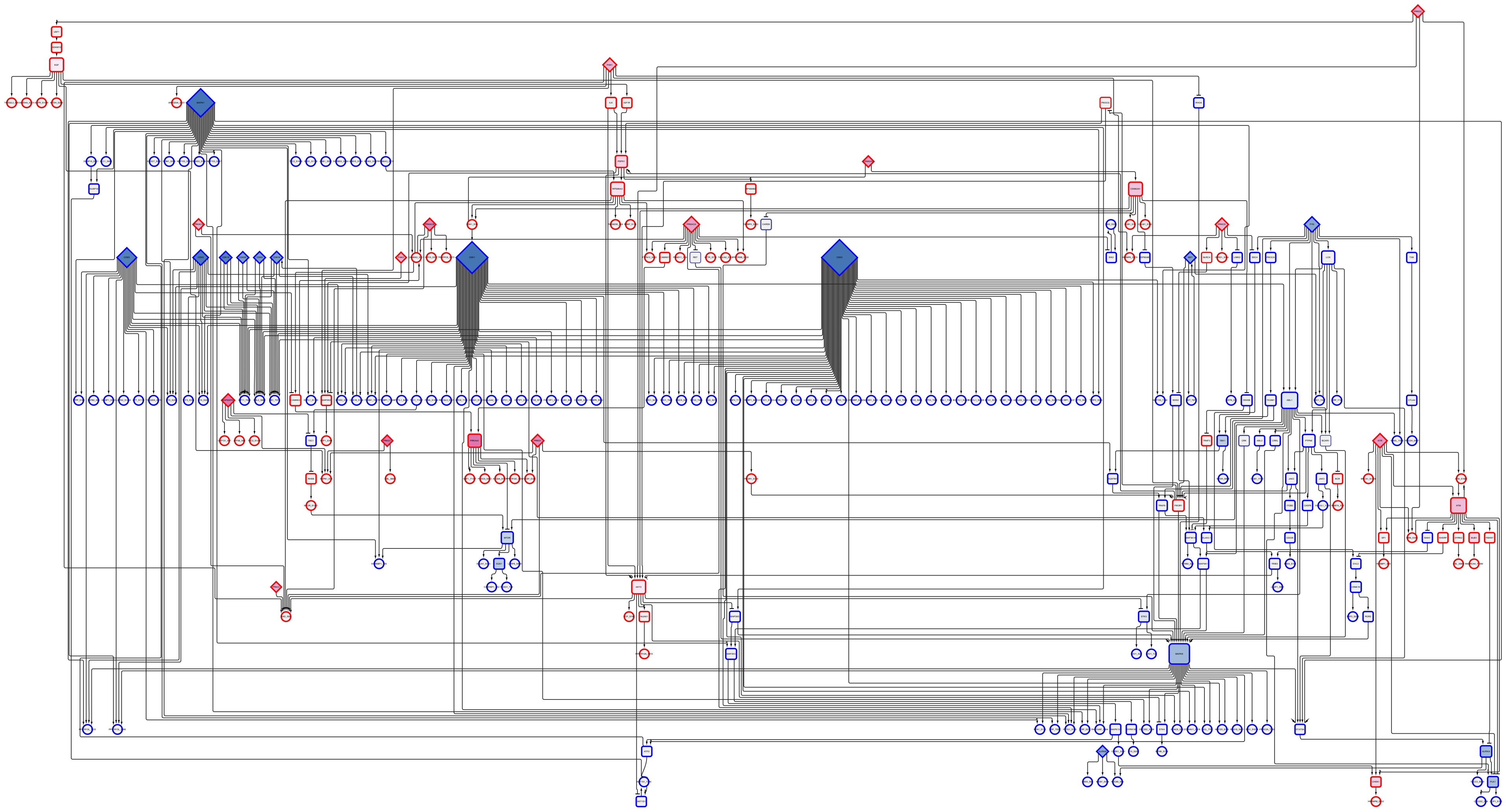

Supplementary Figure 6K

# SW948

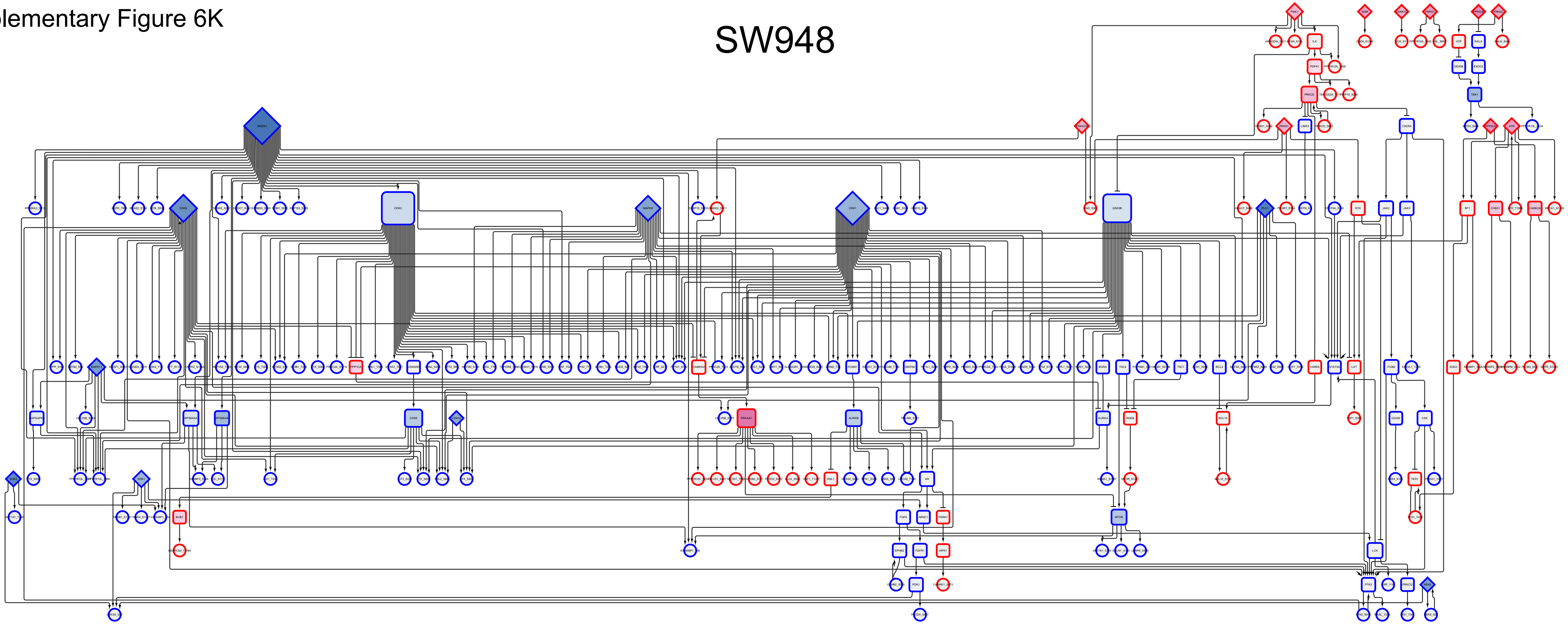

Supplementary Figure 6L

# T84

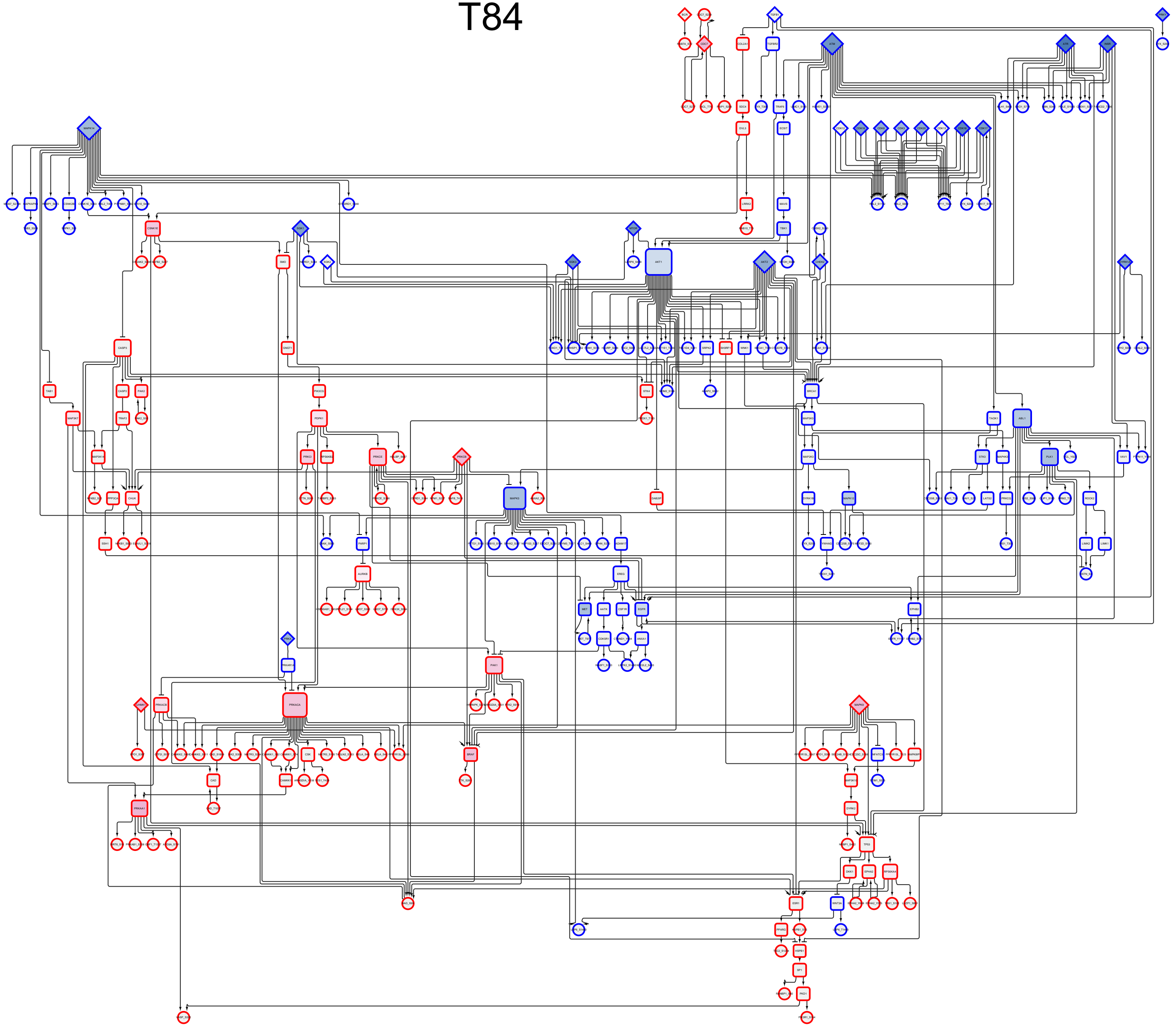

### Supplementary figure 7

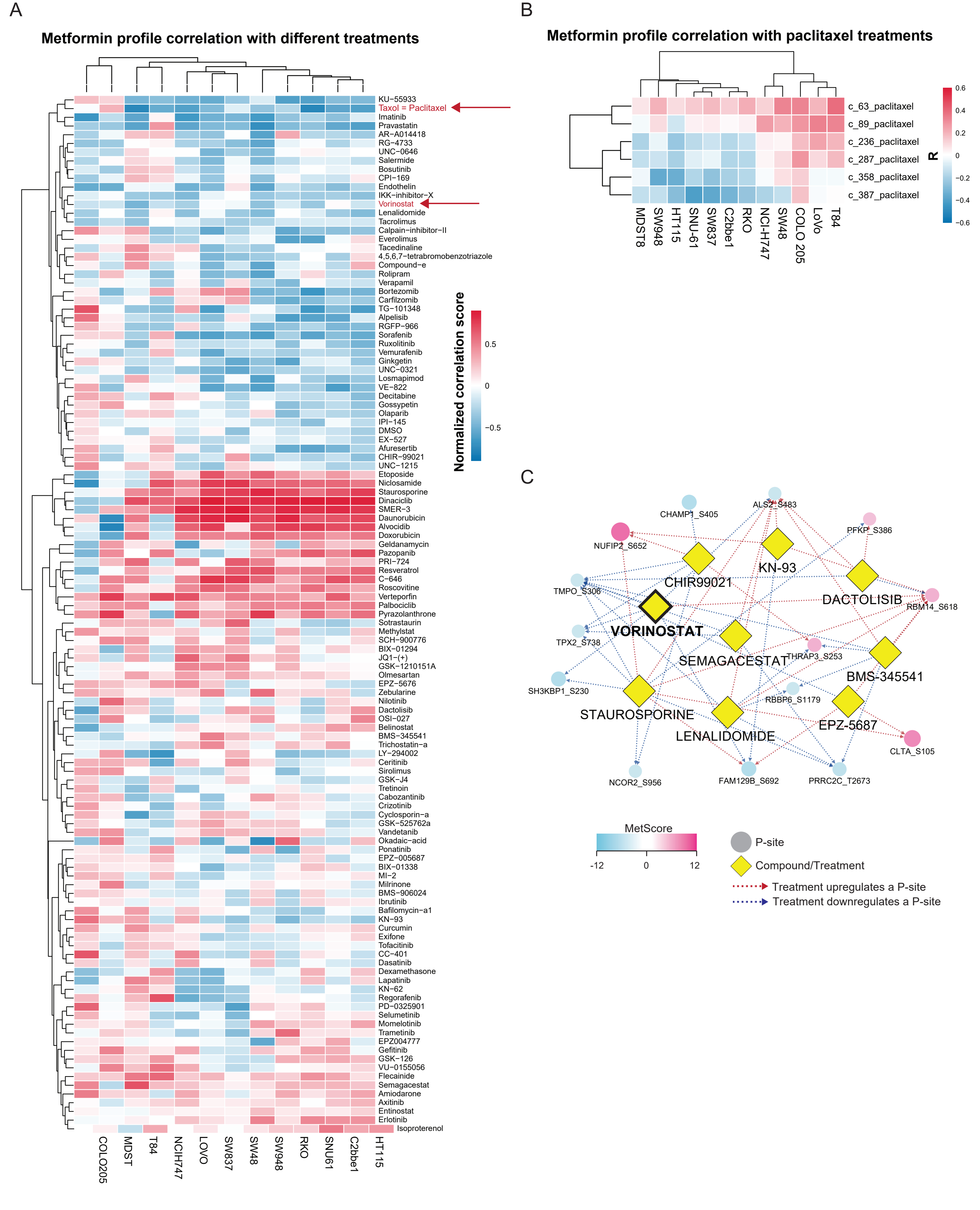
